# Functional importance and paralog-specificity of the intrinsically disordered RsmE C-terminus region in *Pseudomonas fluorescens* Pf0-1

**DOI:** 10.64898/2026.09.16.752055

**Authors:** Meghan K. Wells, McKenna L. Carroll, Peter Fazio, Mayelin Ebersole, Lea Kasmer, Sarah Cole, Anton F. Evans, Wook Kim

## Abstract

RsmE and its homologs function as RNA-binding post-transcriptional regulators that control the production of diverse secondary metabolites. Pseudomonads harbor varying numbers of RsmE paralogs that are often generalized to overlap in function. However, recent studies indicate that these paralogs bind to both overlapping and unique sets of mRNA. In *Pseudomonas fluorescens* Pf0-1, RsmE exclusively regulates the production of multiple extracellular secretions, distinguishing it from its paralogs, RsmA and RsmI. While the majority of the three paralogs’ core sequences are highly conserved, each possesses a vastly different C-terminus region. The C-terminus tails are generalized to be functionless, as they are intrinsically disordered and devoid of known mRNA binding sites. Here, we challenge this notion through analyses of various naturally emergent RsmE variants that differentially impact repressive function solely from changes in the C-terminus tail. Engineered Rsm chimeras containing the various paralog cores and tails further confirmed the functional importance of the C-terminus tail, as neither replacing the RsmE tail with the RsmA tail nor attaching the RsmE tail to a different paralog core restored repression. However, the RsmI tail complemented function in the presence of the RsmE core, indicating that RsmE’s functional specificity lies within the core despite the essentiality of the C-terminus. Bioinformatic analyses also revealed that the C-terminus tail composition of RsmA to be the most unique among the paralogs and each tail sequence is uniquely and highly conserved across diverse proteins, not only in bacteria, but also in eukaryotic species.

**IMPORTANCE:** The post-transcriptional regulator RsmE controls the production of various extracellular secretions in *Pseudomonas fluorescens* Pf0-1, making it functionally unique from its paralogs, RsmA and RsmI. The three paralogs share highly conserved sequences and secondary structures save for their disordered C-termini, which are generalized to be functionless. Here, we demonstrate that the absence of the RsmE C-terminus tail completely abolishes native function and that various alterations to the tail differentially impact repression of extracellular secretions. The results described here strongly support functional uniqueness of Rsm paralogs, challenge the supposed dispensability of their C-terminus tails, and deepen our understanding of RsmE-regulation, which modulates biofilm formation and virulence in clinically significant bacteria.

## INTRODUCTION

In densely structured microbial communities, the capacity to dominate space and nutrients determines whether individual genotypes thrive or perish. The post-transcriptional regulator RsmE plays a key role in spatiogenetic structure formation within *Pseudomonas fluorescens* colonies by modulating the production of multiple extracellular secretions (1–3). RsmE belongs to the CsrA/RsmA (carbon storage regulator/regulator of secondary metabolism) family, homologs of which regulate diverse cellular processes by sequestering a broad range of mRNA (4–6). CsrA/RsmA proteins primarily function by binding to a GGA motif overlapping the Shine Dalgarno sequence, thus blocking ribosome binding and preventing the translation of cognate mRNA (7–18). CsrA/RsmA homologs regulate diverse social and virulence traits across the Gammaproteobacteria (5, 19).

*Pseudomonas* species typically possess two to four Rsm paralogs, all of which carry highly conserved residues that dictate homodimerization and RNA-binding (20). Although Rsm paralogs have been described to be functionally redundant, such generalization appears to be context dependent. Previous work exploring their role in intercellular interactions in *P. fluorescens* Pf0-1 revealed that RsmE is functionally unique from its two paralogs, RsmA and RsmI. Isolated mucoid patches naturally emerge within densely populated wild type (WT) colonies specifically due to mutations in *rsmE*, consequently resulting in the de-repression of multiple extracellular products (1). When the native function of RsmE is either fully or partially abolished, the respective mutant forms a continually expanding isogenic spatial structure against the surrounding WT cells, with strikingly reduced cellular density. These extracellular secretions include a polysaccharide that pushes away the neighboring WT cells, a biosurfactant that spatially localizes other secretions to maintain the genotypic boundary, and a type VI secretion system which kills WT cells that invade the expanding patch (2, 3). Together, these secretions allow *rsmE* mutants to dominate over the neighboring cells by creating and protecting optimal space within a crowded WT population. In contrast, neither *rsmA* nor *rsmI* mutations are observed in the emergent patches. Production of both the extracellular polysaccharide and biosurfactant remains unaffected in Δ*rsmA* or Δ*rsmI*, and mucoid patches naturally emerge within these mutant colonies that are phenotypically identical to those formed by naturally occurring *rsmE* mutants in a WT population (2).

CsrA/Rsm proteins typically comprise five β-sheets, an α-helix, and an unstructured C-terminus tail (7, 8, 20, 21). They form a homodimer with antiparallel β-sheets creating a hydrophobic core between the two sheets and the α-helix along with two protruding C-terminus tails (7, 9, 20), to simultaneously bind two separate mRNA targets (8, 9, 21). Despite minor sequence and structural differences, Rsm paralogs possess very similar stabilizing contacts when in complex with mRNA targets (21), contributing to the general notion that Rsm paralogs have overlapping or cumulative regulatory functions (11, 21–26). In *P. fluorescens* Pf0-1, all three paralogs are expressed in WT and have highly conserved primary sequence and secondary structure, yet, only the deletion of *rsmE* leads to spatial structure formation and increased relative fitness over the WT (2). Then, what makes RsmE functionally unique from its paralogs?

Rsm paralog sequences in *P. fluorescens* can be arbitrarily classified into two distinct regions (Figure 1): the core (first ∼50 residues), which is highly conserved and responsible for folding, dimerization, and mRNA binding, and the C-terminus tail (last 10-13 residues), which is highly variable across both paralogs and orthologs (27). The β-sheets and α-helix are known to be essential for dimerization and mRNA binding. However, the absence of secondary structure and considerable sequence divergence at the C-terminus of CsrA/Rsm homologs have led to the generalization that it does not impact dimerization or RNA binding, and therefore holds no function (20, 21, 27). All known residues that are important for RNA binding reside within the core, with majority of sites being in the first and fifth β-sheets (9, 28). In *P. fluorescens* specifically, the solution structure of RsmE has been resolved, and the C-terminus tail was not shown to be directly involved in mRNA binding, at least for one of its known targets (9).

**Figure 1.**
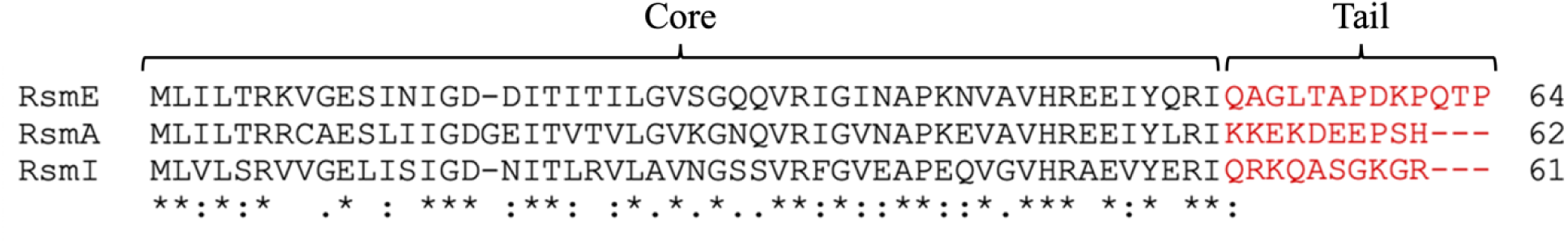
Multiple sequence alignment of the three Rsm paralogs in *Pseudomonas fluorescens* Pf0-1. The core (black) is highly conserved, while the C-terminus tail (red) is highly variable between paralogs. Multiple sequence alignment was created using Clustal Omega (64). Residue positions are marked based on conservation, indicating fully conserved positions (*), strong conservation (:), weak conservation (.), or no conservation (no symbol).

We had previously built a library containing over 500 *rsmE* mutants that naturally and independently emerged as an isolated patch within WT *P. fluorescens* Pf0-1 colonies, which includes 12 unique mutations within or just upstream of the C-terminus tail of RsmE (1). We thus hypothesize that the C-terminus tail is not only involved in, but vital to RsmE’s repressive function. Here, we explore the functional importance and specificity of RsmE’s C-terminus tail by characterizing the impact of the naturally derived mutations and engineered chimeric Rsm paralogs on extracellular secretions and competition against the WT.

## RESULTS

### Alterations in the RsmE C-terminus tail differentially impact extracellular polysaccharide (EPS) and biosurfactant production

To assess the importance of the C-terminus tail in RsmE’s native repressive function, 12 previously isolated *rsmE* mutants were analyzed for EPS and biosurfactant production, each harboring a unique mutation within or just upstream from the C-terminus tail (Figure 2A). These mutants arose naturally from WT colonies as an independent patch and were previously classified as either red or blue based on their ability to produce the biosurfactant following one day of incubation: red mutants produce it and blue mutants do not (1). In contrast, all mutants produce the extracellular polysaccharide (EPS) leading to the mucoid colony morphology (1), and reconfirmed here for all tail mutants (Figure 2B). Biosurfactant production was also reassessed for each tail mutant, but over multiple days this time, along with WT and Δ*rsmE* serving as controls. Although the five red mutants and seven blue mutants behaved as previously observed on day one (1), a small ring was visible for five blue mutants on day two (Figure 2C), and no ring was detected for the remaining two blue mutants even after three days (Figure 2D). Contrary to our initial red/blue categorization (1), biosurfactant production does not appear to be binary, but rather manifests in varying degrees depending on the mutation.

**Figure 2.**
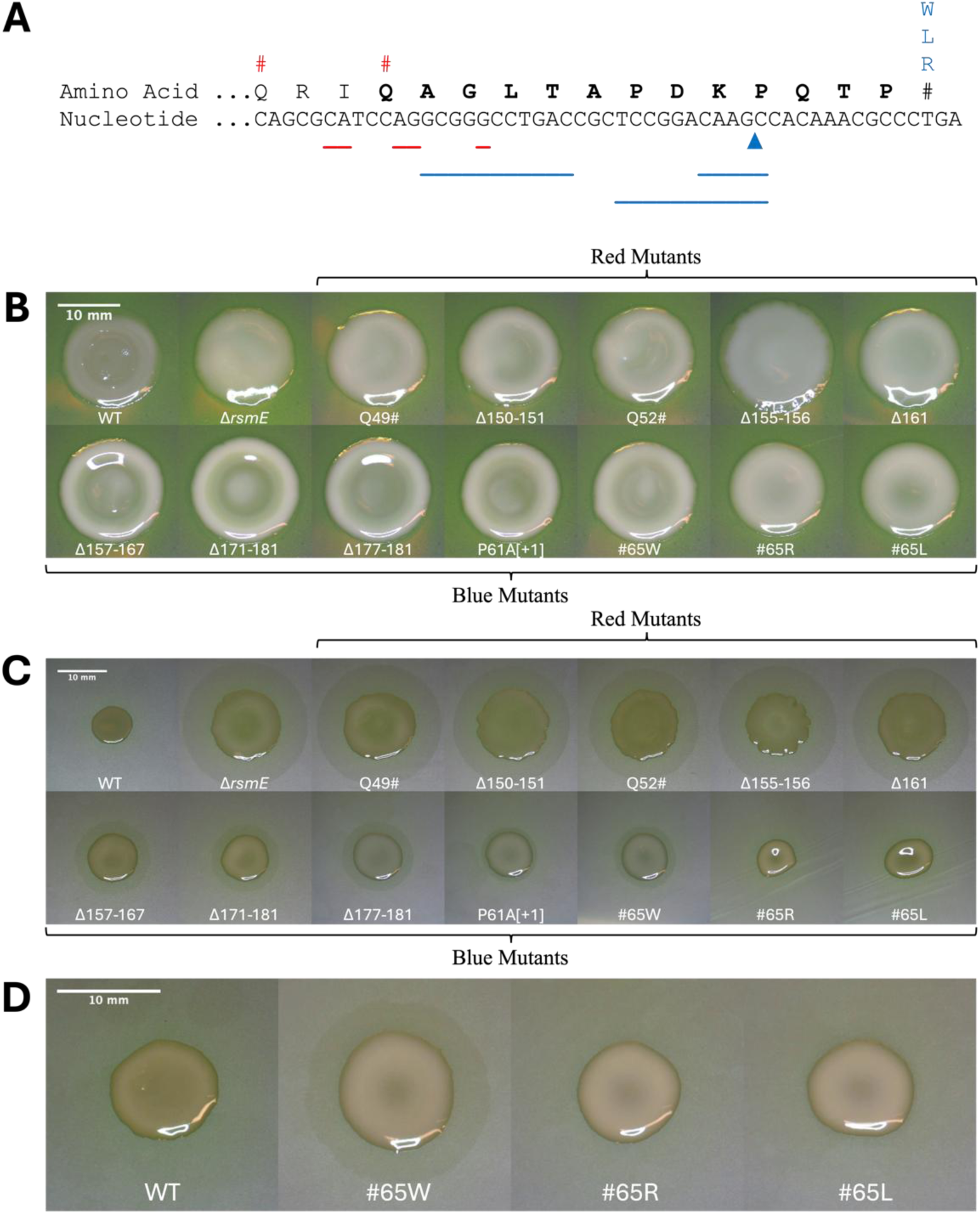
Naturally derived C-terminus tail *rsmE* mutants universally produce the EPS but produce varying amounts of the biosurfactant. (A) Locations of the twelve different mutations within or near the C-terminus tail of *rsmE*. Lines indicate deletions, arrows indicate insertions, hashtags indicate nonsense mutations, and letters indicate nonstop mutations. Mutants are color coded based on their initial classification of red (surfactant producing) or blue (not surfactant producing) after one day of growth. Bolded residues dictate the C-terminus tail. (B) Morphology of the *rsmE* tail mutants after three days of growth on PAF. (C) Surfactant production of *rsmE* tail mutants represented by production of a surfactant ring around the colony when grown on the dull side of a polycarbonate membrane overlain on PAF. Images were captured after two days of growth. (D) Surfactant assay of WT and the three stop-loss mutants after three days of growth. Mutants #65R and #65L are the only mutants that do not produce a surfactant ring like the WT. Scale bars represent 10 mm.

We next quantified biosurfactant production by measuring the diameter of the ring at day two, which revealed three clear categories (Figure 3A). Group 1 displayed production comparable to Δ*rsmE*, group 2 produced a moderate amount compared to Δ*rsmE*, while group 3 produced no biosurfactant like the WT. A clear pattern emerges in reference to the primary sequence, where mutations closer to the beginning of the C-terminus tail (i.e. closer to the core) exert more impact on the repressive function of RsmE (Figure 3B). Δ161 and #65W appear to be outliers, which are explored further in the following section. In general, the more the original C-terminus tail remains, the better the altered RsmE retains its repressive function on biosurfactant production. In contrast, the entire C-terminus tail region of RsmE appears to be essential for repressing EPS production, which plays a crucial role in space expansion within a crowded population (2).

**Figure 3.**
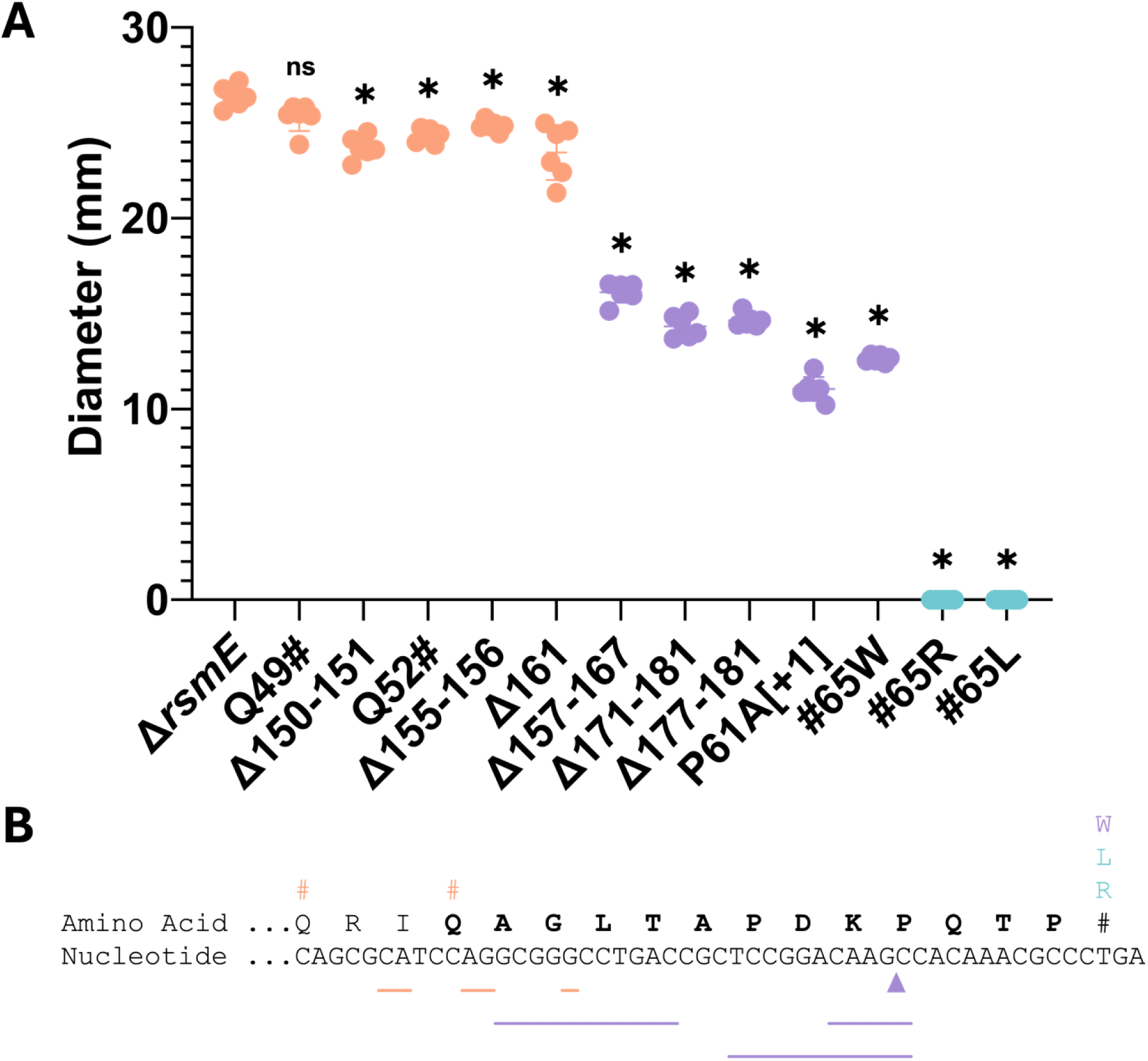
New classification of the blue tail mutants. (A) Quantification of biosurfactant production of the C-terminus tail *rsmE* mutants reveals three clear groups. The diameter of the surfactant ring was measured after two days of growth. Comparing the surfactant measurements across mutants (n = 6) revealed a significant difference (ANOVA; *P* < 0.0001). Pairwise comparisons were made using Tukey’s honestly significant difference test (*P* < 0.05), and significance is shown relative to Δ*rsmE* (denoted by *) or noted as nonsignificant (ns). While post-hoc pairwise comparisons indicate significant difference of most mutants due to low experimental variance, the macro-level separation of the three color-coded phenotypic clusters represents a clear biological effect. (B) New categories of natural tail mutants indicate that alterations closer to the core have more of an impact on repressive function. Lines indicate deletions, arrows indicate insertions, hashtags indicate nonsense mutations, and letters indicate nonstop mutations. Mutants are color coded based on their level of surfactant production on day two, with substantial (orange, group 1), moderate (purple, group 2), or no (turquoise, group 3) biosurfactant produced. Bolded residues dictate the C-terminus tail.

### *rsmE* tail mutations have varying impact on predicted primary and secondary protein structure

To explore why the tail mutations cause varying impact on biosurfactant production, bioinformatic analysis was carried out to predict consequences of each mutation on the primary and secondary protein structures (Figure 4). Among the group 1 mutations, which result in substantial biosurfactant production comparable to Δ*rsmE*, three result in a complete or premature truncation of the C-terminus tail. The other two mutations in group 1 result in a frameshift, adding bulk (∼75 additional residues after site of mutation) to the tail of the protein, and the change occurs far enough upstream that all original C-terminus tail information is lost due to the mutation. The large drop in biosurfactant production from group 1 to group 2 suggests that mutations that are further downstream from the core-tail junction result in significantly more retention of RsmE’s native function. One apparent outlier in group 1 is Δ161, which physically clusters with group 2 mutations in terms of the relative distance from the junction. However, the Δ161 mutation results in an immediate stop codon associated with the frameshift, which should cause a severe defect in function like the other upstream nonsense mutations that truncate the tail.

**Figure 4.**
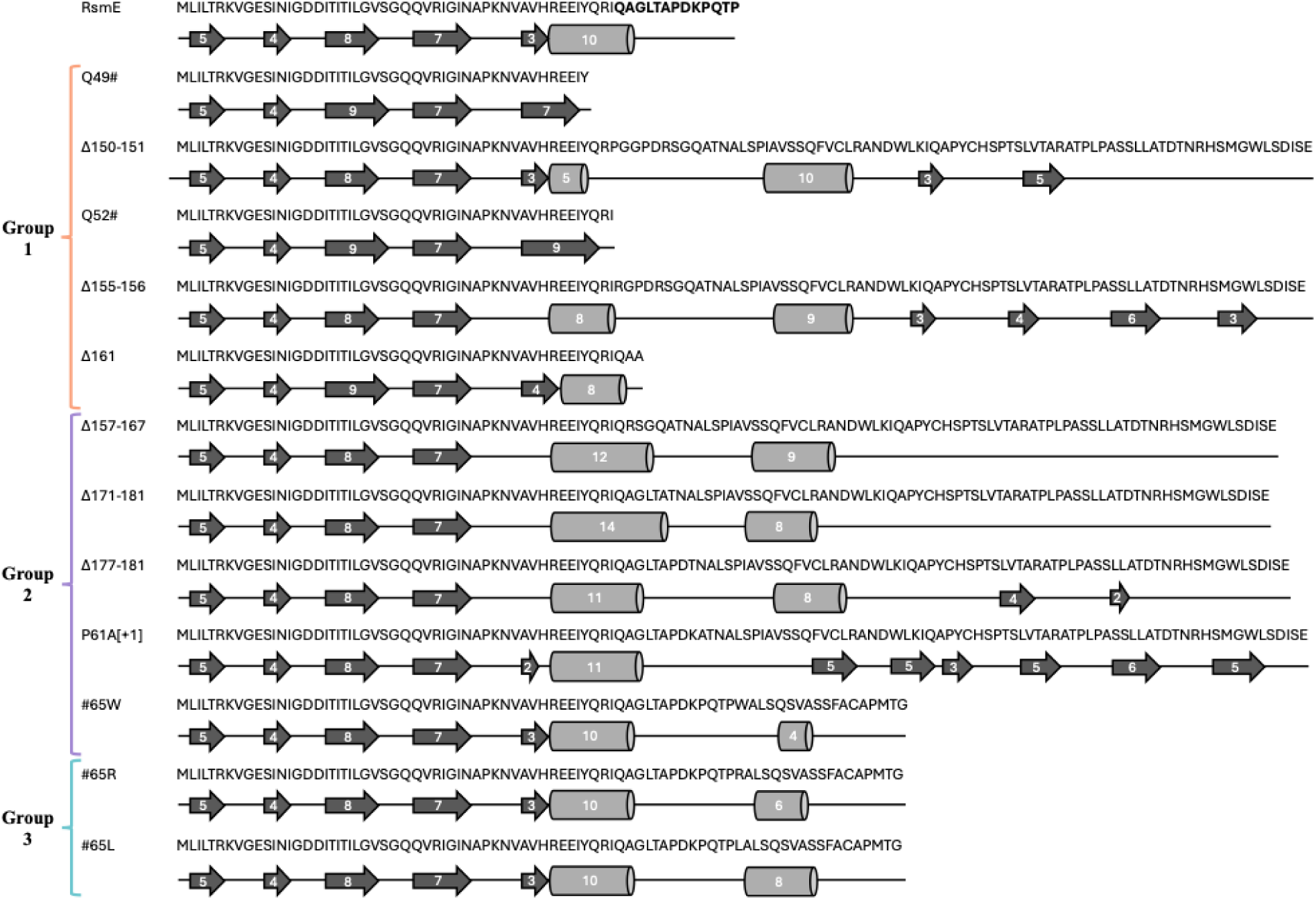
C-terminus tail *rsmE* mutations differentially impact primary and secondary structures. Arrows represent β-sheets and cylinders represent α-helices. The numbers inside indicate the number of amino acids involved in each indicated structure. Bolded residues in the RsmE sequence represent the C-terminus tail. All secondary structure predictions were made using Phyre2.2 (63). Mutants are color coded based on their ability to produce substantial (Group 1; orange), moderate (Group 2; purple), or no (Group 3; turquoise) biosurfactant.

Group 2 mutations also add significant bulk (66-71 additional residues after the site of mutation) to the tail of the protein caused by a frameshift (Figure 4). However, due to the mutations occurring further downstream compared to the group 1 mutations, some of the C-terminus tail information is retained in the group 2 mutants. This pattern supports the notion of retaining more of the native function based on the relative distance from the core-tail junction. The group 3 mutations that do not produce any biosurfactant are stop-loss mutations that add relatively moderate bulk (18 residues) to the tail, but the original C-terminus tail information entirely remains (Figure 4). The respective mutants retain full function in terms of repressing biosurfactant production, but the associated changes in the tail length appears to have some functional impact since the repression of the EPS is abolished like all tail mutants (Figure 2B). The apparent outlier is the stop-loss mutation #65W, which falls in group 2 for producing a moderate amount of the biosurfactant. While tryptophan is the largest and most hydrophobic amino acid, it is unclear why the #65W mutation specifically causes greater loss of function when all three stop-loss mutations should result in an identical read through (Figure 4).

Nevertheless, the length of additional bulk at the C-terminus and less retention of the original C-terminus tail information both appear to correlate with the severity of functional loss. Changes in the C-terminus sequence do not appear to impact secondary structures in the core region, with the exception of the fifth β-sheet in certain cases. Although some mutations are predicted to either destabilize or alter the length of the fifth β-sheet, there appears to be no discernible pattern across groups 1 and 2.

### Fitness advantage over the WT is not quantitatively dependent on biosurfactant production

We next asked how the observed differences in the quantity of biosurfactant production affects each mutant’s ability to outcompete the WT. When RsmE is rendered entirely nonfunctional (e.g. red mutations or Δ*rsmE*), all natively repressed extracellular products consequently become de-repressed in the respective mutant, resulting in their spatial dominance over the WT in co-culture (1, 2). Additionally, when biosurfactant production is genetically knocked out in Δ*rsmE*, the relative fitness of the respective mutant over the WT decreases compared to that of Δ*rsmE* (2). Here, we competed each C-terminus tail mutant against WT to assess changes in their relative fitness (Figure 5). A relative fitness value greater than one indicates that the mutant outcompetes the WT, while a value equal to one indicates the two competing isolates are equally fit. As expected, group 1 mutants, which produce comparable amounts of the biosurfactant as Δ*rsmE*, also generated comparable relative fitness values as Δ*rsmE*. Group 3 mutants, which produce no biosurfactant, generated significantly reduced relative fitness values compared to Δ*rsmE*, and mirrored the magnitude of reduction as that of the biosurfactant knockout in Δ*rsmE* (2). However, group 2 mutants, which produce biosurfactant at decreased levels compared to Δ*rsmE* (Figure 3A), yielded relative fitness values not significantly different from that of Δ*rsmE* (Figure 5). These data indicate that while certain tail mutations differentially impact the quantity of biosurfactant production, any production is sufficient to maintain a significant fitness advantage over the WT.

**Figure 5.**
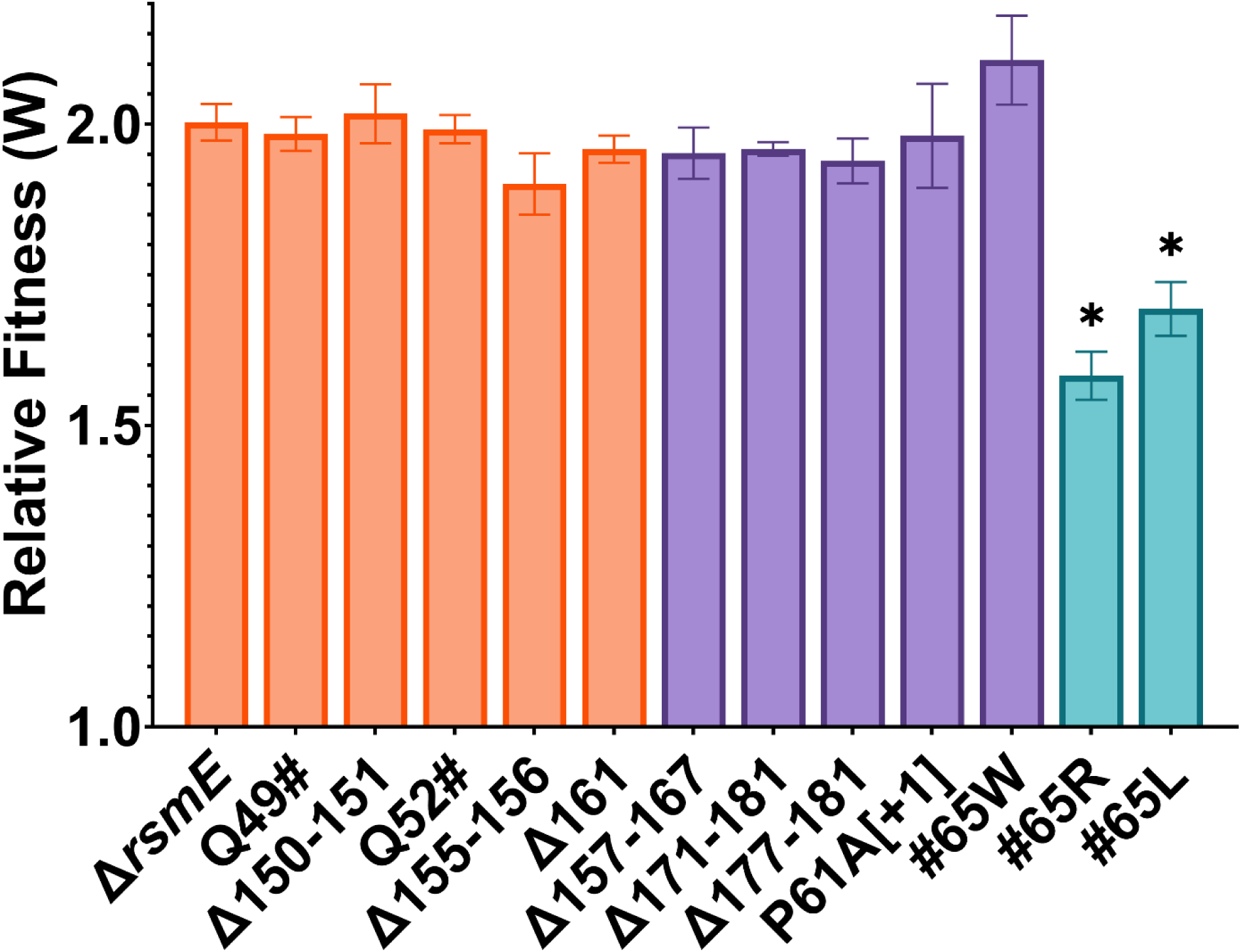
Relative fitness values of the C-terminus tail mutants versus WT. Isolates were competed against WT for four days, then relative fitness was calculated. A relative fitness value (W) of one indicates that the mutant and WT are equal in fitness, while W > 1 indicates that the mutant outcompeted the WT. Comparing the relative fitness values across all competitions revealed a significant difference (ANOVA; *P* < 0.0001). Significance relative to the Δ*rsmE*-WT control is displayed on the graph (denoted by *; Tukey’s HSD; *P* < 0.05). Groups one (orange) and group 2 (purple) mutants produce surfactant and display relative fitness values similar to the Δ*rsmE* control, while group 3 (turquoise) mutants, that do not produce any surfactant, show a significant decrease in relative fitness compared to Δ*rsmE*.

### The specific sequence of the C-terminus tail of RsmE is not essential for repressing biosurfactant production

Given the varying impact of the naturally derived mutations on biosurfactant production, we next explored whether the C-terminus tail of RsmE confers functional specificity from its paralogs. A set of chimeric constructs was engineered to encode the core of RsmE, including the native promoter and 5’ untranslated regions of *rsmE*, and the C-terminus tails of each of the three paralogs (Figure 1). Each construct was then integrated as a single copy into the same non-coding region of the Δ*rsmE* chromosome and screened for restoration of biosurfactant repression. As expected, the chimera with the RsmE core and tail (RsmE/E) fully restored repressive function, not producing a ring like the WT (Figure 6A). However, the chimera with the RsmE core and RsmA tail (RsmE/A) produced a very small ring (Figure 6A), which became more visible after an additional day of growth (Figure 6B). RsmE/A is phenotypically comparable to that of the naturally derived group 2 mutants (Figure 2C), implying partial recovery of function. In contrast, the chimera with the RsmE core and RsmI tail (RsmE/I) did not produce a ring like RsmE/E (Figure 6A), even at day 3 (Figure 6B), indicating full restoration of repressive function. The C-terminus tail of RsmE is clearly essential for function, since biosurfactant production is fully de-repressed in group 1 mutants (Figure 3A). However, the native sequence of the RsmE tail appears to be relatively less important, since the RsmI tail could functionally replace it. Interestingly, RsmA and RsmI tails are equal in length and shorter than that of RsmE (Figure 1), indicating that the mere presence of a C-terminus tail of comparable length is insufficient for maintaining full function.

**Figure 6.**
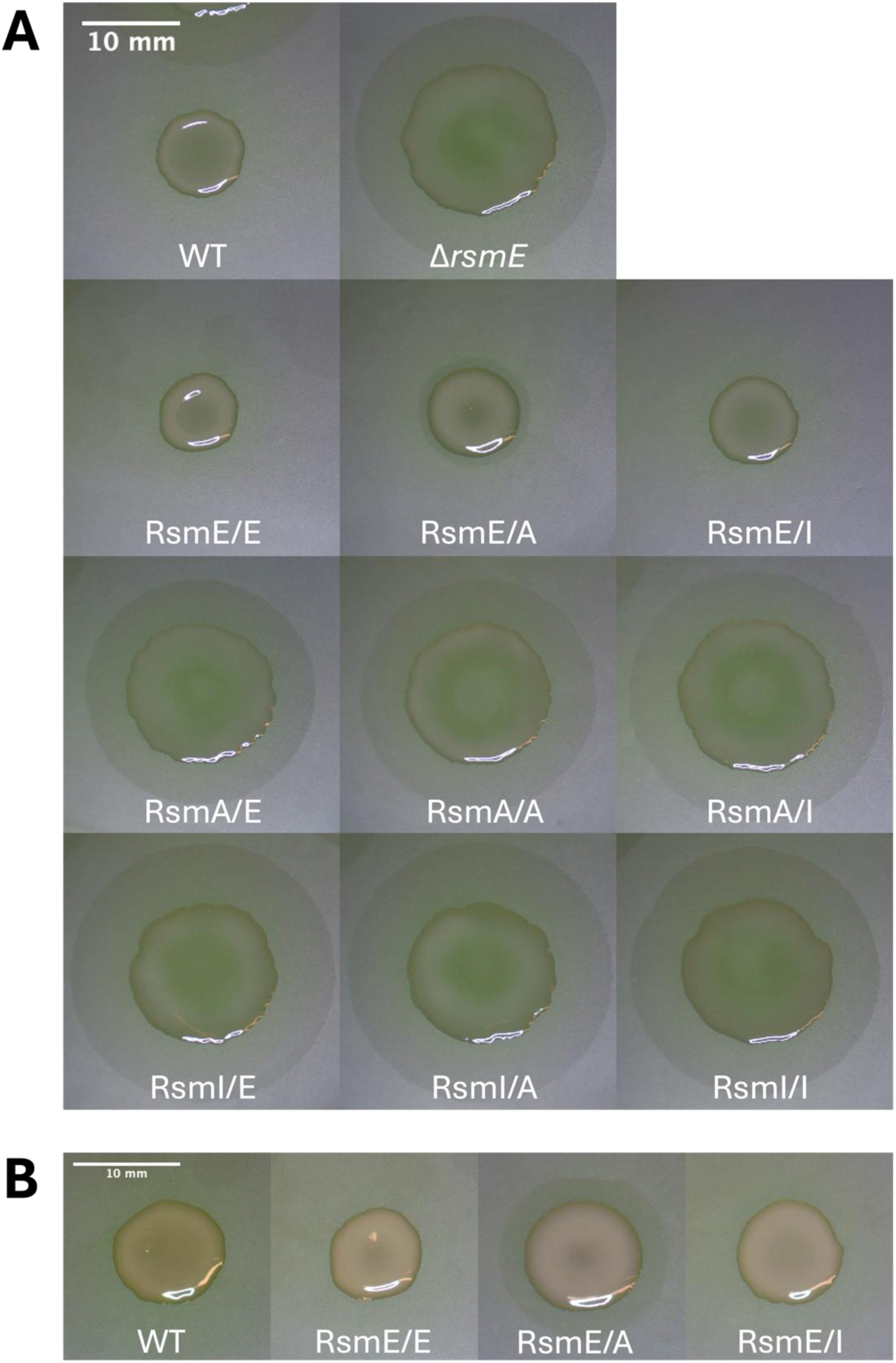
Rsm chimeras produce varying levels of the biosurfactant. Images were taken after two (A) and three (B) days of growth on a polycarbonate membrane overlain on PAF. Absence of a surfactant ring indicates that the given Rsm chimera has restored repressive function of the surfactant in Δ*rsmE*. RsmE/E and RsmE/I are the only chimeras that completely restore repressive function. RsmE/A partially restores repressive function, and none of the RsmA- or RsmI-core chimeras restore repressive function. Scale bars represent 10 mm.

### The core region of RsmE specifies unique function among the paralogs

In contrast to RsmE, neither RsmI nor RsmA natively represses biosurfactant or EPS production (2), yet the RsmE C-terminus tail is functionally essential (Figure 2, Figure 3A) and also appears to be functionally equivalent to that of RsmI (Figure 6). Then, the functional specificity of RsmE from its paralogs must primarily reside in the core region. To address this hypothesis, additional chimeras were constructed and integrated as a single copy in the chromosome of Δ*rsmE*, comprising the RsmA core with the three paralog tails (RsmA/E, RsmA/A, and RsmA/I), and the RsmI core with the paralog tails (RsmI/E, RsmI/A, and RsmI/I). All six chimeras produced the biosurfactant in comparable amounts as Δ*rsmE* (Figure 6A), indicating that simply adding the RsmE tail to the core of either paralog fails to restore the repressive function.

As an independent evaluation for the source of RsmE’s functional specificity from its paralogs, the colony morphology of all nine chimeras was compared, which specifically addresses EPS production (2). Only the RsmE/E and RsmE/I chimeras exhibited repression of EPS production like the WT (Figure 7). RsmE/A appears to be phenotypically similar to Δ*rsmE*, which contrasts from the significant reduction in biosurfactant production (Figure 6). An additional phenotypic difference observed among the chimeras is the emergence of isolated patches (Figure 7). As described previously, these patches naturally emerge from WT colonies due to mutations in *rsmE* (1, 2), including the C-terminus tail mutations described above (Figure 2A). When *rsmE* is knocked out, the entire colony is mucoid, and no patches emerged (Figure 7). Consistent with the absence of the mucoid phenotype, isolated patches emerged only in WT, RsmE/E, and RsmE/I colonies, further confirming that the RsmI C-terminus tail is functionally equivalent to that of the RsmE. Whole genome sequencing confirmed that a select emergent patch in both RsmE/E and RsmE/I was caused by a unique nonsense mutation within the respective core region of the chimeric gene (Table S1). All RsmA- and RsmI-core chimeras are mucoid and did not produce patches (Figure 7).

**Figure 7.**
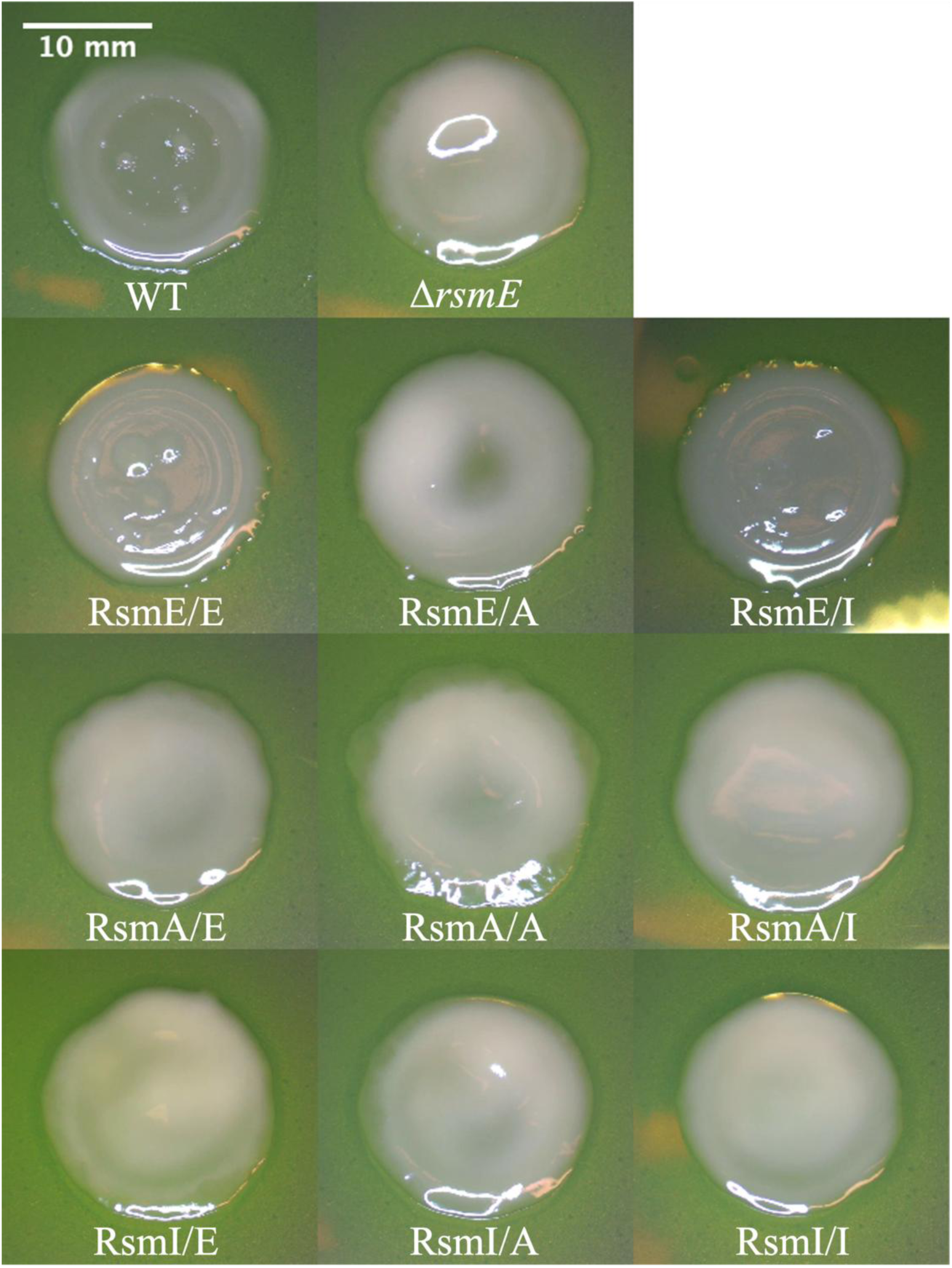
RsmE/I is functionally equivalent to both RsmE/E and native RsmE in repressing EPS production. Chimeras were grown on PAF and imaged after four days of growth. RsmE/E and RsmE/I are morphologically identical to WT. Mucoid patches naturally emerge only in WT, RsmE/E, and RsmE/I colonies. Scale bar represents 10 mm.

The same pattern was observed in the competitions of the Rsm chimeras against the WT (Figure 8). RsmE/E and RsmE/I produced relative fitness values comparable to that of the WT vs WT competition, indicating that both chimeras are equally fit as the WT. Conversely, RsmE/A and all of the RsmA- and RsmI-core chimeras generated relative fitness values that are not significantly different from that of Δ*rsmE* (Figure 8). Taken together, the core region of RsmE appears to primarily dictate the functional specificity from its paralogs, while the C-terminus tail provides a secondary, but essential role with less sequence rigidity.

**Figure 8.**
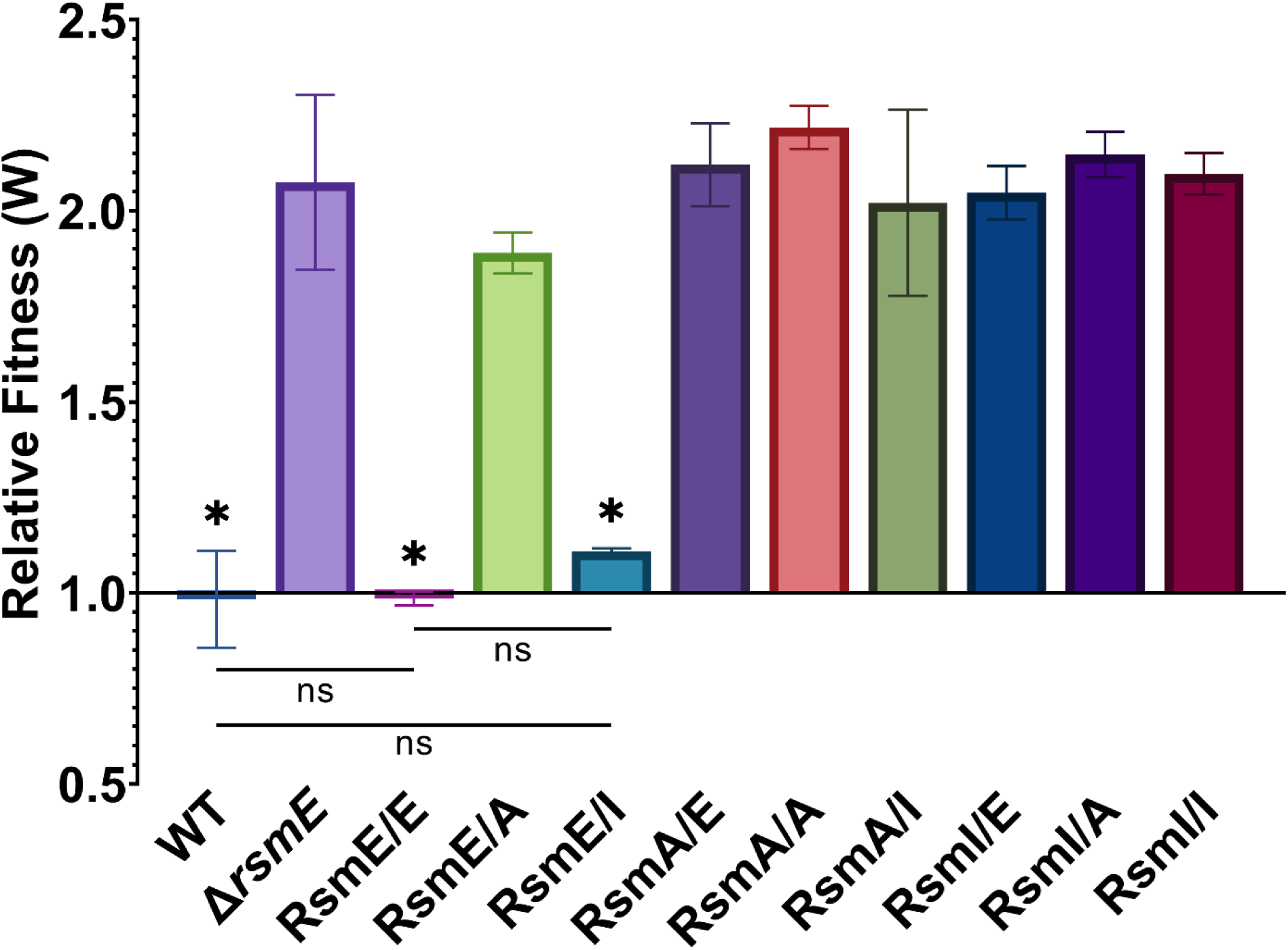
The chimeras do not impact the relative fitness over the WT with the exception of RsmE/E and RsmE/I. A relative fitness value (*W*) equal to one indicates that the competing isolate is as equally fit as the WT in co-culture colonies. *W* > 1 indicates the isolate outcompetes the WT. *W* values significantly different from the Δ*rsmE* versus WT competition are noted by the asterisks. Significance was determined using a one-way ANOVA (*P* < 0.0001) followed by Tukey’s honestly significant difference test (*P* < 0.05) for pairwise comparisons. Chimera competitions significantly different from the Δ*rsmE*-WT control (denoted by *) and not significantly different from the WT-WT control (denoted by ns) are shown. The comparison between RsmE/E and RsmE/I competitions is also noted as not significant. RsmE/E and RsmE/I have similar fitness to WT, while RsmE/A and the RsmA- and RsmI-core chimeras have similar fitness to Δ*rsmE*.

### Residue properties vary among paralog C-terminus tails

Because the C-terminus tail of RsmE is vital for function but not fully dependent on primary sequence, we next sought to assess each Rsm paralog tail sequence for any unique patterns in amino acid property composition by measuring variations in R-group charge, size, and hydrophobicity (Figure 9). The RsmA tail contains a high percentage of charged residues (80%), with an equal number of alternating positively and negatively charged residues that result in a net charge of zero. The RsmE tail also has a neutral net charge but is comprised of fewer charged residues (∼15%), while the RsmI tail lies in between its two paralogs in quantity of charged residues (40%). However, the RsmI tail stands apart as the only strongly cationic sequence, exhibiting complete depletion of negative charges and a highly positive net charge (+4). Tail residues were also grouped by size based on the volume of their R-group as small (<60 Å^3^) or large (≥60 Å^3^). The RsmA tail contains mostly large (80%) and few small residues (20%). The RsmE and RsmI tails show a more even distribution, containing 46% and 40% small residues, respectively. Hydrophobicity of each amino acid R-group was classified using the Kyte-Doolittle hydrophobicity scale, which considers not only charge and polarity but also the residue’s tendency to preside on the hydrophilic surface or hydrophobic core of a folded protein The RsmE and RsmI tails contain some hydrophobic residues, 23% and 10% respectively. Conversely, the RsmA tail is completely hydrophilic. Although the RsmA tail stands apart from its paralogs due to its high quantity of charged residues, large size, and lack of hydrophobicity, it remains unclear what role, if any, these different properties play in functional specificity among the paralogs.

**Figure 9.**
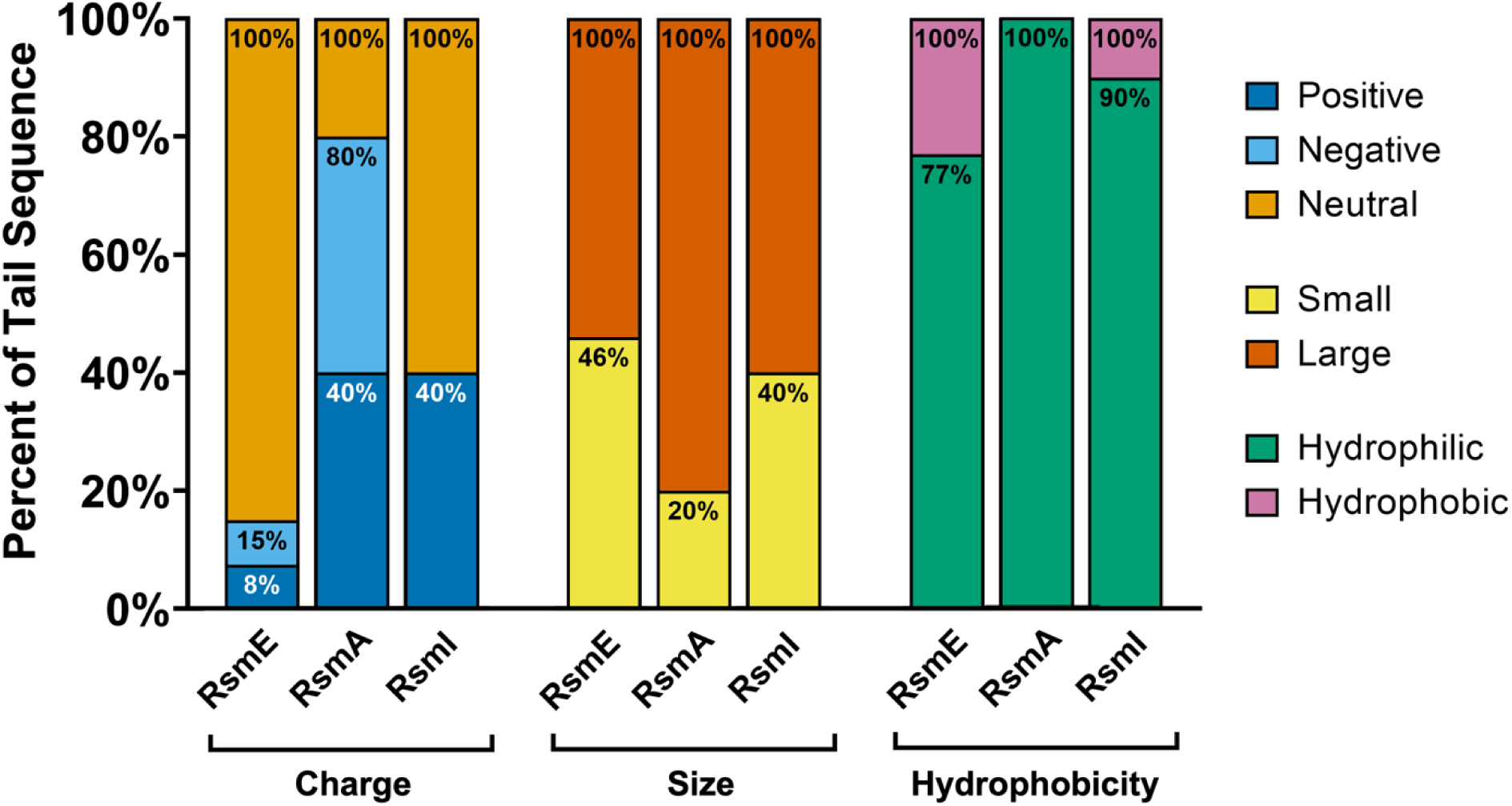
Residue property distribution of Rsm paralogs show differences in charge, size, and hydrophobicity. Each residue in the paralogs’ tail sequences were sorted based on various amino acid characteristics. Residue charge was categorized based on the R-group as positively charged, negatively charged, or neutral. Size was sorted by the volume of the R-group into small (<60 Å^3^) and large (≥60 Å^3^). Hydrophobicity of each amino acid R-group was classified using the Kyte-Doolittle hydrophobicity scale (29).

### C-terminus tails of each Rsm paralog are highly conserved across diverse species

To assess the evolutionary conservation of the Rsm paralog C-terminus tail sequences, we conducted short-sequence optimized Blastp and filtered the top 1,000 hits for 90% query coverage and 90% identity (Table S3). The filtered results were analyzed for presence across various kingdoms, *Pseudomonas* groups, and annotated functions. A summary of these results is displayed in Table 1. *Pseudomonas* groups were categorized using the NCBI taxonomy browser Functional annotations were grouped using the Clusters of Orthologous Genes (COG) database (31), and the Evolutionary Genealogy of Genes: Non-supervised Orthologous Groups (EggNOG) database (32, 33) was used to assign COG codes to eukaryotic proteins.

**Table 1.** Categorizations of paralog tail Blastp hits with 90% coverage and identity.

| Description | RsmE | RsmA | RsmI |
| --- | --- | --- | --- |
| Panel A: Represented Kingdoms |  |  |  |
| Bacteria | 31 | 74 | 44 |
| *Pseudomonas* | 29 | 58 | 18 |
| *Other/General* | 2 | 16 | 26 |
| Archaea | 0 | 1 | 0 |
| Plant | 0 | 4 | 0 |
| Animal | 0 | 4 | 8 |
| Fungi | 0 | 6 | 1 |
| Protist | 0 | 4 | 1 |
| Panel B: *Pseudomonas* Groups<sup>a</sup> |  |  |  |
| *P. fluorescens* | 4 | 9 | 4 |
| *P. chlororaphis* | 0 | 2 | 0 |
| *P. syringae* | 0 | 11 | 0 |
| *P. aeruginosa* | 0 | 1 | 0 |
| *P. putida* | 0 | 1 | 0 |
| Unclassified | 25 | 34 | 14 |
| Non-*Pseudomonas* | 2 | 35 | 36 |
| Panel C: COG Categories<sup>b</sup> |  |  |  |
| 1. Information Storage & Processing | 0 | 7 | 4 |
| 2. Cellular Processes & Signaling | 31 | 68 | 41 |
| *CsrA Annotated* | 31 | 65 | 18 |
| *Non-CsrA Annotated* | 0 | 3 | 23 |
| 3. Metabolism | 0 | 1 | 1 |
| 4. Poorly Characterized | 0 | 17 | 8 |

<sup>a</sup> *Pseudomonas* groups categorized using the NCBI taxonomy browser (30).
<sup>b</sup> Annotated functions categorized using the Clusters of Orthologous Genes (COG) database (31).
Eukaryotic proteins were assigned COG codes using the Evolutionary Genealogy of Genes:
Non-supervised Orthologous Groups (EggNOG) database (32, 33).

Application of the stringent filter parameters resulted in 31 hits for the RsmE tail, 93 for RsmA, and 54 for RsmI, and each paralog search yielded a unique set of hits. The RsmE tail search produced not only the least hits, but also the least variation, representing only bacteria. Of the bacteria, 29 were *Pseudomonas* species within only the *P. fluorescens* group, and all hits were also annotated as CsrA. Thus, the RsmE tail appears highly conserved only in CsrA family proteins in closely related *Pseudomonas* species. In contrast, the RsmA tail search yielded the most diversity across kingdoms and *Pseudomonas* species, which is expected since RsmA is considered the ancestral paralog (27). The hits ranged across all kingdoms, including 74 bacteria, 1 archaea, 4 plants, 4 animals, 6 fungi, and 4 protists (Table 1). Of the 74 bacteria, 58 are *Pseudomonas* species that span 5 different *Pseudomonas* groups, confirming broad conservation among *Pseudomonads*. CsrA annotations account for 65 of the hits, nearly 70%, while the remainder are distributed among the three other COG categories of information storage and processing, metabolism, and poorly characterized.

The RsmI tail search fell between its paralogs in diversity, representing 44 bacteria, 8 animals, 1 fungus, and 1 protist (Table 1). However, only 18 of the 54 hits were *Pseudomonas* species, all of which fell into the *P. fluorescens* group or were unclassified. Interestingly, while most of the hits fell into the COG category of cellular processes and signaling, CsrA annotations comprise only about 33% of the category. The RsmI tail sequence appears to be specialized within *Pseudomonas* species but also shows conservation in a wide variety of proteins outside of carbon storage regulators. Additionally, for all three paralogs, conserved sequences were not limited to the respective C-terminus tail region. Although short length sequences may be randomly preserved, 100% conservation of contiguous amino acid residues across all domains of life implies that they do hold functional significance, further challenging the common notion that the Rsm paralog C-terminus tails are functionless.

## DISCUSSION

Our combined study of the naturally emergent C-terminus tail mutants and the engineered chimeras strongly indicates that the functional specificity of RsmE from its two paralogs is driven by its core residues, that are already known to determine folding, dimerization, and mRNA binding (9, 28). We also demonstrate the importance of the C-terminus tail of RsmE, which has long been assumed to be functionless. The C-terminus tail is necessary for RsmE-mediated repression of extracellular secretion production, but it does not appear to define RsmE’s functional specificity from its paralogs.

Analysis of the naturally emergent RsmE tail mutants identified a time-dependent appearance of the biosurfactant, as different alterations to the C-terminus differentially impacted repressive function. Repression of the EPS is much more tail sequence-rigid, as none of the naturally derived tail mutants retained the ability to repress EPS production. However, the RsmI tail of the chimeric RsmE/I protein fully restored repression of both secretions, indicating that the RsmI tail somehow functionally parallels the RsmE tail. Although the specific primary sequence of the C-terminus tail does not appear to be absolutely critical, the amino acid content still matters, since the RsmA tail, which is equivalent in length to the RsmI tail and shorter than the RsmE tail, significantly impacts biosurfactant production. Several of the naturally emergent *rsmE* tail mutants add bulk to the C-terminus, which may indirectly alter mRNA binding by physically blocking the active sites. A recent study has shown that the RsmE-mediated EPS and biosurfactant production, along with a newly characterized type VI secretion system (T6SS), are all indirectly regulated by RsmE, as deletion of *rsmE* alters both the transcription of the respective genes and protein levels (3). This pattern also holds true in other species, as RsmE homologs have been found to bind hundreds of mRNA targets, including diverse transcription regulators and signal transduction systems (4, 6, 34–36).

The C-terminus tail of RsmE does not appear to participate in either the tertiary or quaternary structure of the homodimer, since it was resolved without defined presence of the tail region (9). Although the cognate mRNA that co-resolved with the homodimer, *hcnA*, does not depend on the tail for interaction with the core, it remains possible that other mRNA interactions could variably rely on the tail for stabilization. Since NMR spectroscopy, X-ray crystallography, and cryo-EM rely on structural averaging, the constant motion of disordered, flexible protein termini often renders these dynamic regions uninterpretable in final structural models. However, static structure is not a prerequisite for biological function. Intrinsically disordered protein regions (IDRs) frequently utilize their conformational plasticity to allow for diversity in function or multifunctionality (37–39). Intrinsically disordered C-termini specifically have been shown to be involved in a variety of functions across all domains of life, including, but not limited to, tuning binding affinity to and selection of binding partners, targeting for degradation, chaperoning folding, and facilitating turnover rates of binding partners (40–42).

Consistent with other documented IDRs, the three Rsm paralog C-terminus tails in *P. fluorescens* are enriched in disorder-promoting residues, which are often small, charged, polar, and/or structure-disrupting (e.g. A, G, R, T, S, K, Q, E, and P), and depleted in bulky, hydrophobic residues (38, 43–45). Distinctive properties of the native paralog tails include the highly charged and completely hydrophilic RsmA tail, the proline-rich RsmE tail, and the strongly cationic RsmI tail. Highly charged IDRs have been shown to drive liquid-liquid phase separation (LLPS) through weak multivalent contacts with disorder-promoting residues (44, 45), which can lead to differential partitioning of proteins and their binding partners depending on what is recruited to the condensate (46, 47). For example, the disordered C-terminal region of Ribonuclease E (RNase E) in *Caulobacter crescentus*, which consists of clustered blocks of alternating positive and negative charges like RsmA, creates condensates called bacterial ribonucleoprotein-bodies (BR-bodies) through LLPS that provide localized sites for RNA degradation (46). A role of the C-terminus tails in localization of the paralogs within the cytoplasm could result in different binding partners among the paralogs.

Unlike RsmA, the RsmI tail lacks negatively charged residues, resulting in a positive net charge (+4). Enrichment of positively charged residues (and depletion of negative charges) in IDRs is a conserved and valuable trait in ribosomal proteins to attract the negatively charged RNA phosphate backbone (43). IDRs in ribosomal proteins also contribute to a variety of additional functions, with the most dominant being protein-protein interactions (43). The possibility of recruiting accessory proteins to the Csr/Rsm-RNA complex remains, as the majority of research efforts are directed toward the RNA interactome, leaving very little experimental evidence of direct or indirect protein interactors of Csr/Rsm homologs (27). Two direct protein-binding partners are known to regulate CsrA in distantly related species: CesT, through competitive inhibition (48), and FliW, through allosteric inhibition (49–52).

Interestingly, the extended C-terminus tail of CsrA is required for FliW antagonism (52–54), but species that evolved to regulate Csr/Rsm proteins through sRNAs instead of FliW, like *P. fluorescens*, lost both the proteinaceous antagonist and the extended C-terminal domain of CsrA that allows protein binding (27, 52).

The proline-rich C-terminus tail of RsmE potentially retains a propensity for protein binding, as proline-rich IDRs often form a secondary structure called polyproline II (PPII) helices, which are highly abundant in intrinsically disordered proteins (55, 56) and commonly involved in protein-protein interactions (57–59). While PPII helices can be detected in experimentally solved or modeled structures, they are often overlooked in standard analyses (56), reinforcing the plausibility that the RsmE C-terminus tail may mediate protein binding.

Additionally, the RsmE and RsmI C-terminus tails contain hydrophobic residues that can drive protein-protein interactions through hydrophobic contacts, while the RsmA tail is completely hydrophilic. Although this difference could explain why the RsmE/I chimera retains repressive function while RsmE/A does not, it does not provide a reason for RsmE/A’s partial recovery of repression.

A recent study of RsmE in *Pseudomonas protegens* may justify RsmE/A’s phenotype. Finol et al. demonstrated that the disordered C-terminus of RsmE mediates a conformational switch between two semi-*holo* intermediate states, in which one RNA is bound to the dimer (60). This exchange allosterically alters the binding affinity of the second site, facilitating the transfer of bound mRNA targets to regulatory sRNAs to relieve repression. Replacing the RsmE C-terminus tail with the highly charged RsmA tail may reduce binding affinity via unfavorable electrostatic interactions, thereby enabling the ribosome to outcompete the RsmE/A chimera and initiate intermittent translation. In contrast, the strongly cationic C-terminus tail of RsmE/I may restore repressive function by attracting the negatively charged backbone of RNA targets, stabilizing protein-RNA contacts and increasing overall binding affinity.

Additionally, interactions between the C-terminus tail residues A57 and P58 and the bound RNA and RsmE α-helix, respectively, were determined to be vital to the slowly exchanging semi-*holo* states and thermodynamically coupled to the empty binding site (60). Interestingly, in our group 1 naturally derived *rsmE* tail mutants, which completely relieve repression of the biosurfactant, residues A57 and P58 are either both altered or entirely missing, while the group 2 and group 3 mutants, which differentially produce biosurfactant, retain both residues, with the exception of two group 2 mutants (Δ157-167 and Δ171-181) that contain a P58T substitution. Thus, the role of the disordered C-terminus tail of RsmE described by Finol et al. is consistent with the observations described here.

Together, our analyses of the naturally emergent *rsmE* C-terminus tail mutants and Rsm chimeras demonstrate that RsmE’s functional specificity lies in the core, yet its disordered C-terminus tail is vital for repression of the EPS and biosurfactant. While the explicit mechanism remains unclear, the data presented here unmistakably demonstrate the importance of the disordered RsmE C-termini, refuting the generalized notion that they are functionless. The collection of naturally derived RsmE tail mutants and Rsm chimeras also provide a unique route to further studying functional specificity among the paralogs and the importance of disordered regions in protein function.

## METHODS

### Strains and culture conditions

Lennox LB medium (Fisher) was used for both solid and liquid overnight cultures. *Pseudomonas* agar F (PAF; Difco) was used for phenotype screens. *Pseudomonas* minimum medium (PMM; 3.5 mM potassium phosphate dibasic trihydrate, 2.2 mM potassium phosphate monobasic, 0.8 mM ammonium sulfate, 100 mM magnesium sulfate, 100 mM sodium succinate) was used to isolate *P. fluorescens* strains from *E. coli* strains. Routine cloning used *E. coli* JM109 (Promega), and S17.1λpir (61) was used as the donor strain in conjugations. All *E. coli* strains were incubated at 37 °C, and *P. fluorescens* strains were incubated at either 30 °C or room temperature (∼22 °C). All liquid cultures were incubated with shaking at 250 rpm. When required, antibiotics were added at the following final concentrations: ampicillin (100 μg/mL), streptomycin (50 μg/mL), kanamycin (50 μg/mL), and chloramphenicol (6 μg/mL). All *Pseudomonas* isolates used in this study are listed in Table S1.

### Construction of antibiotic-resistant isolates

All natural RsmE C-terminus tail mutants were tagged with a kanamycin resistance marker via the mini-Tn7 chromosomal insertion system as described previously (1, 62). All conjugation mixtures included the host *P. fluorescens* mutant, the donor strain containing the antibiotic resistance marker (pUC18TminiTn7T-Km in S17.1λpir *E. coli*), a helper strain containing the transposase gene (pUX-BF13 in S17.1λpir *E. coli*), and a mobilizer strain containing the mobile elements required for conjugation (pRK600 in HB101 *E. coli*). All four strains were rinsed in 1X PBS, mixed in equal parts, and spotted onto an LB plate. The conjugation mixture was incubated at 30 °C overnight. After incubation, the mixture was scraped off the LB plate, resuspended in PBS, serially diluted, and plated onto PMM-Km to select for the tagged *P. fluorescens* mutants. Chromosomal insertion of the antibiotic resistance marker was also confirmed through PCR (5’-ATGGGATCGGCCATTGAACAAG and 5’-GAAGAACTCGTCAAGAAGGCGATA). Both the kanamycin and streptomycin resistance markers have been demonstrated to be neutral for relative fitness measurement competition experiments in *P. fluorescens* Pf0-1 (1).

### Construction of Rsm chimeras

Nine chimeric Rsm proteins containing the various cores and tails of each paralog (RsmE/E, RsmE/A, RsmE/I, RsmA/E, RsmA/A, RsmA/I, RsmI/E, RsmI/A, and RsmI/I) were created by first amplifying the *rsm* paralog genes from WT using a forward primer that amplifies starting 500 bp upstream to include necessary promoter regions and reverse primers that contain the end of the core sequence and the entire C-terminus tail sequence. The PCR product was cloned into the pGEMT-Easy (Promega) plasmid then subcloned into the pHRB2 plasmid using the ApaI and SacI restriction enzymes and transformed into S17λpir *E. coli* cells. The chimeric genes were then mated into Δ*rsmE* via conjugation, as described above. All primers used for cloning the chimeras are listed in Table S2.

### Morphology assessment and extracellular polysaccharide detection

To assess morphology, extracellular polysaccharide production, and patch formation of each C-terminus tail mutant and chimera, 20 μL of overnight cell culture was spotted onto PAF plates, incubated at room temperature, and imaged after 3 or 4 days of growth. Each isolate was assessed for mucoidy and patch formation as an indicator of RsmE activity. Formation of patches implies RsmE is functional and repressing secretions, leading to the selection for *rsmE* mutants that naturally emerge due to spontaneous mutation. Conversely, isolates with a nonfunctional RsmE appear mucoid and do not form patches due to secretions already being lifted. Patches formed by RsmE/E and RsmE/I were isolated and whole genome sequenced at SeqCenter (Pittsburgh, PA).

### Biosurfactant assay and quantification

To assess surfactant production of each C-terminus tail mutant and chimera, Nucleopore Track-Etch polycarbonate membranes (Whatman; 0.4 μM pore size, 90-mm diameter) were used as previously described (1, 2). The dull side of the membrane contains gaps and ridges that trap cells but let the biosurfactant permeate through, producing a visible ring around the colony. The membrane was overlain on PAF plates with the dull side of the membrane facing upwards, and 20 μL of cell culture was spotted onto the plates. All surfactant assay plates were incubated at room temperature, and surfactant production was imaged using the Hayear overhead microscope (HY-2307) after two and three days of growth. The diameters of the surfactant rings were measured using Fiji (ImageJ).

### Secondary structure analysis

To produce secondary structure predictions for all natural RsmE C-terminus tail mutants, Phyre2.2 was used (63). Protein sequences were analyzed using normal mode, which uses PSI-Blast to detect sequence homologs, Psi-pred to predict secondary structure, and Diso-pred to predict disorder.

### Relative fitness measurements

Relative fitness was measured by competing the kanamycin tagged mutants and chimeras with streptomycin tagged WT as previously described (1–3). Mutants and chimeras were grown to saturation in LB broth with antibiotic, washed in 1X PBS, then diluted to 10^-3^ in PBS. The diluted kanamycin tagged isolates were mixed 1:1 with undiluted WT suspension. The seed mixture was spotted on PAF plates in three 20 μL spots and incubated at room temperature for four days. To quantify initial population sizes, the competition mixture was serially diluted and spotted on PMM-Km and PMM-Sm plates. Seed plates were incubated at 30 °C overnight. After incubation, colonies were counted, and initial CFU was calculated.

After four days of growth, the competition mixtures were scraped from the PAF plates and resuspended in 5 mL PBS. The mixture was then diluted, spotted onto PMM-Km and PMM-Sm plates, and incubated overnight at 30 °C. The following day, colonies were counted to calculate CFU. Relative fitness of each competing isolate against WT was calculated as follows: [ln(CFU of mutant at time of sampling/CFU of mutant at time zero)]/[ln(CFU of WT at time of sampling/CFU of WT at time zero)].

### Statistical analysis

Each relative fitness competition had three biological replicates and two technical replicates for each biological replicate. The surfactant quantification utilized data from six biological replicates of each mutant after two days of growth. For both the surfactant quantification and competition experiments, the data were first analyzed with a one-way analysis of variance (ANOVA) to assess the significance of the means of the biological replicates.

Tukey’s honestly significant difference test (*P* < 0.05) was then used to make pairwise comparisons within the datasets. All comparisons are noted as statistically significant or nonsignificant. Statistical tests were performed using GraphPad Prism.

### Amino acid property analysis

The amino acid composition of the three Rsm paralog C-terminus tail sequences were analyzed for patterns in charge, size, and hydrophobicity of residue sidechains. Residue charge was categorized based on the R-group as positively charged (R, K, and H), negatively charged (D and E), or neutral (S, T, Q, G, A, L, and P). Size was sorted by the volume of the R-group into small (<60 Å^3^; G, A, S, D, and T) and large (≥60 Å^3^; P, E, H, Q, K, L, and R). Hydrophobicity of each amino acid R-group was classified as hydrophilic (R, K, H, D, E, S, T, Q, G, and P) or hydrophobic (A and L) using the Kyte-Doolittle hydrophobicity scale (29).

### Analysis of homology among Rsm paralog C-terminus tails

Protein Blast (Blastp) searches were performed on the three Rsm protein C-terminus tail sequences (RsmE-QAGLTAPDKPQTP; RsmA-KKEKDEEPSH; RsmI-QRKQASGKGR) to identify conversation of each paralog tail. The search parameters were modified to accommodate the short query sequences, using the NCBI non-redundant (nr) database, the PAM30 scoring matrix, a word size of 2, and an expect value (E-value) of 200,000. The top 1,000 Blastp hits of each paralog tail search were analyzed for the presence of unique genera and annotated functions. The hits were functionally categorized using the Clusters of Orthologous Genes (COG) database (31), and the Evolutionary Genealogy of Genes: Non-supervised Orthologous Groups (EggNOG) database (32, 33) was used to assign COG codes to eukaryotic proteins.

The COG database sorts protein annotations into 4 main and 26 subcategories based on function. The first main category is information storage and processing and includes the subcategories of translation, ribosomal structure, and biogenesis (J), transcription (K), and replication, recombination and repair (L). The second category, cellular processes and signaling, includes signal transduction mechanisms (T), cell wall/membrane/envelope biogenesis (M), cytoskeleton (Z), extracellular structures (W), and post-translational modification, protein turnover, and chaperones (O). The third category is metabolism, which includes lipid transport and metabolism (I) and inorganic ion transport and metabolism (P). The final category is poorly characterized proteins, which includes the subcategories of general function prediction only (R) and function unknown (S).

### Data availability

All noncommercial plasmids or strains used in this study are available for distribution upon request.

## ACKNOWLEDGEMENTS

W.K. designed the study; M.W., M.C., P.F., M.E., L.K., S.C., A.E., and W.K. performed experiments; M.W. and W.K. analyzed data; and M.W. and W.K. wrote the manuscript. This study was funded by the National Institute of General Medical Sciences of the NIH (1R15GM132856 to W.K.).

## SUPPLEMENTARY MATERIAL

**Table S1.**
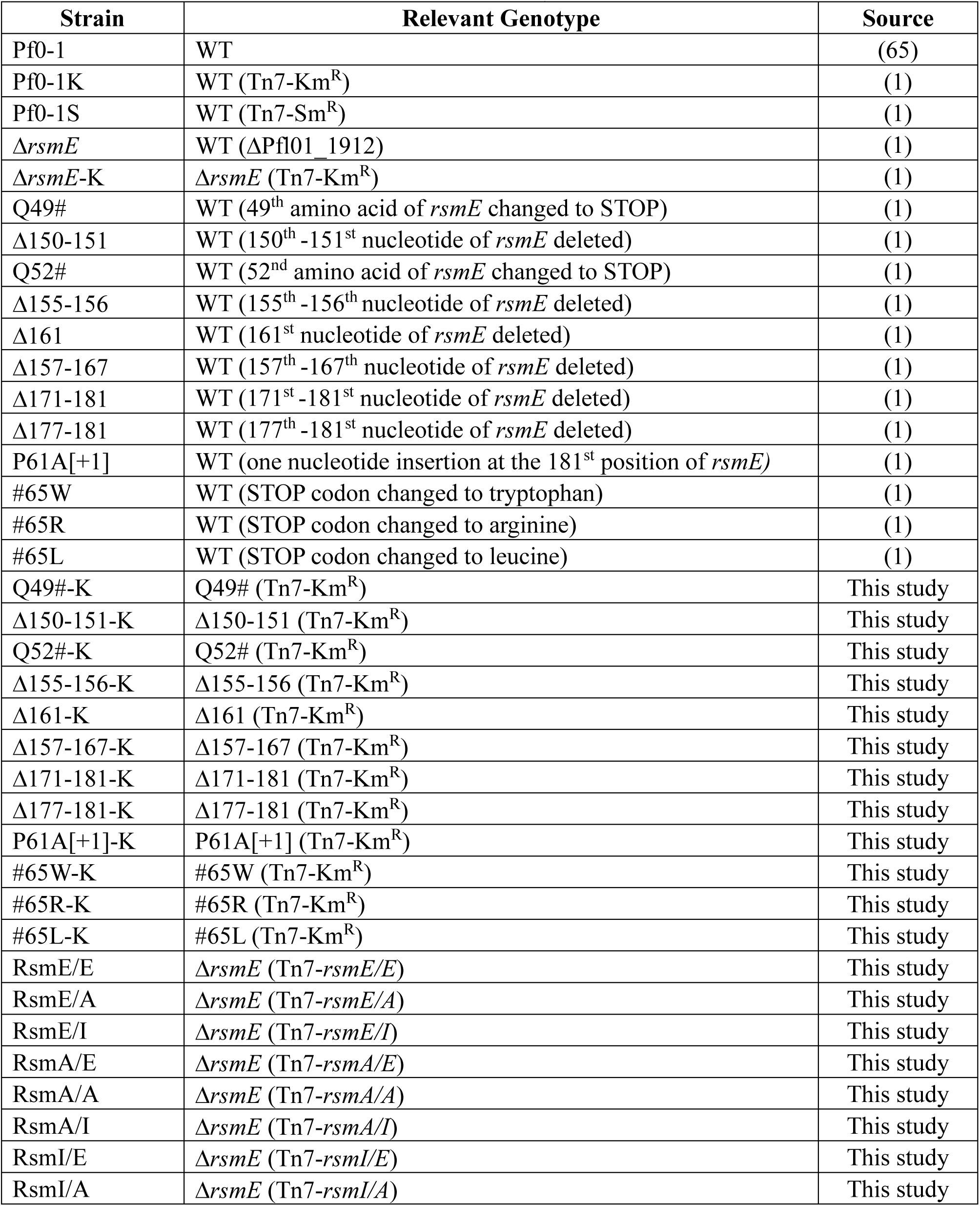
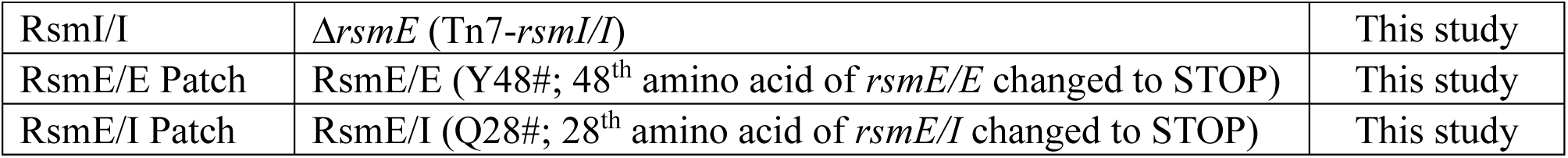
List of *Pseudomonas* strains used in this study.

**Table S2.**
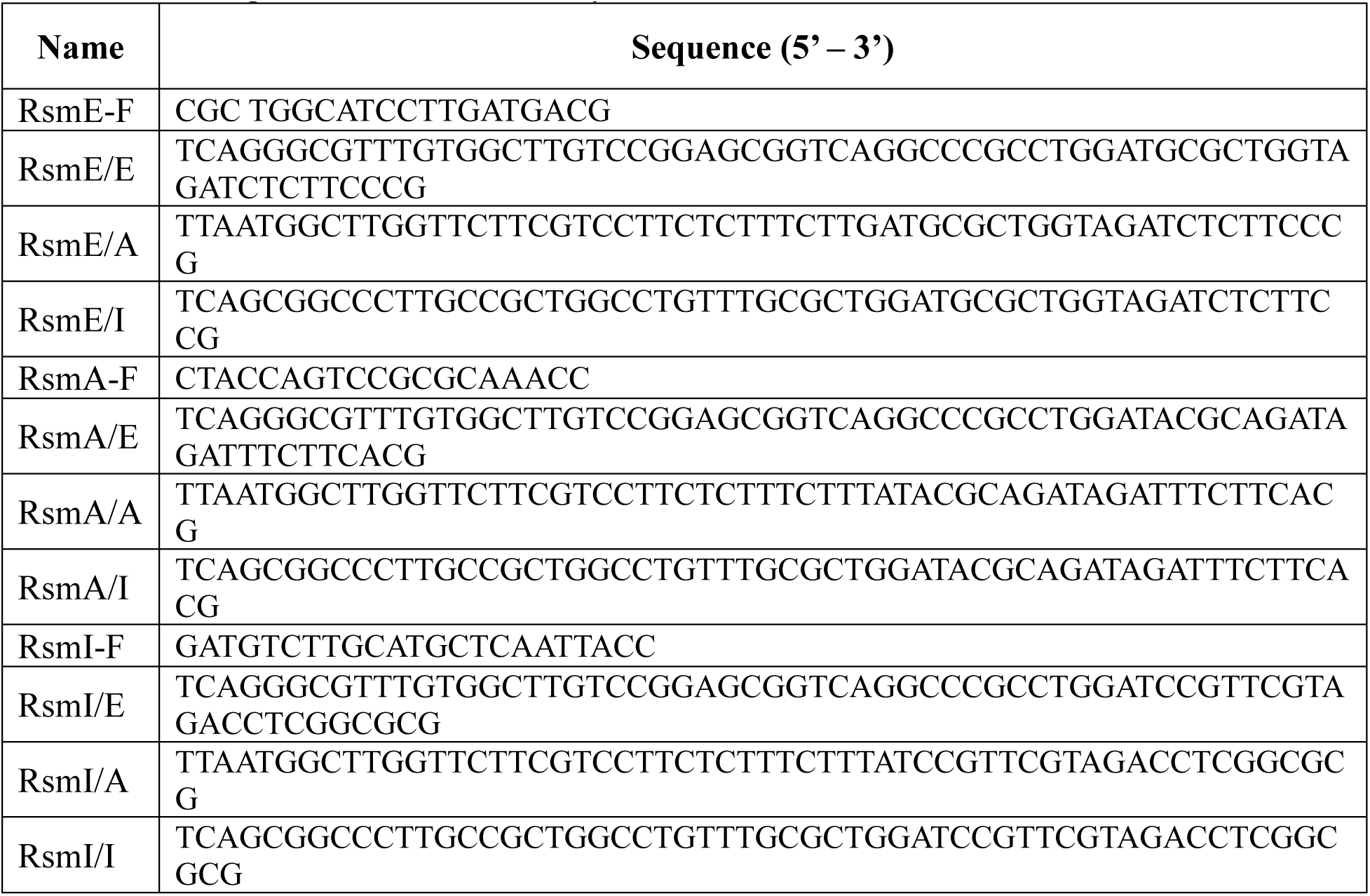
List of primers used in this study.

**Table S3.**
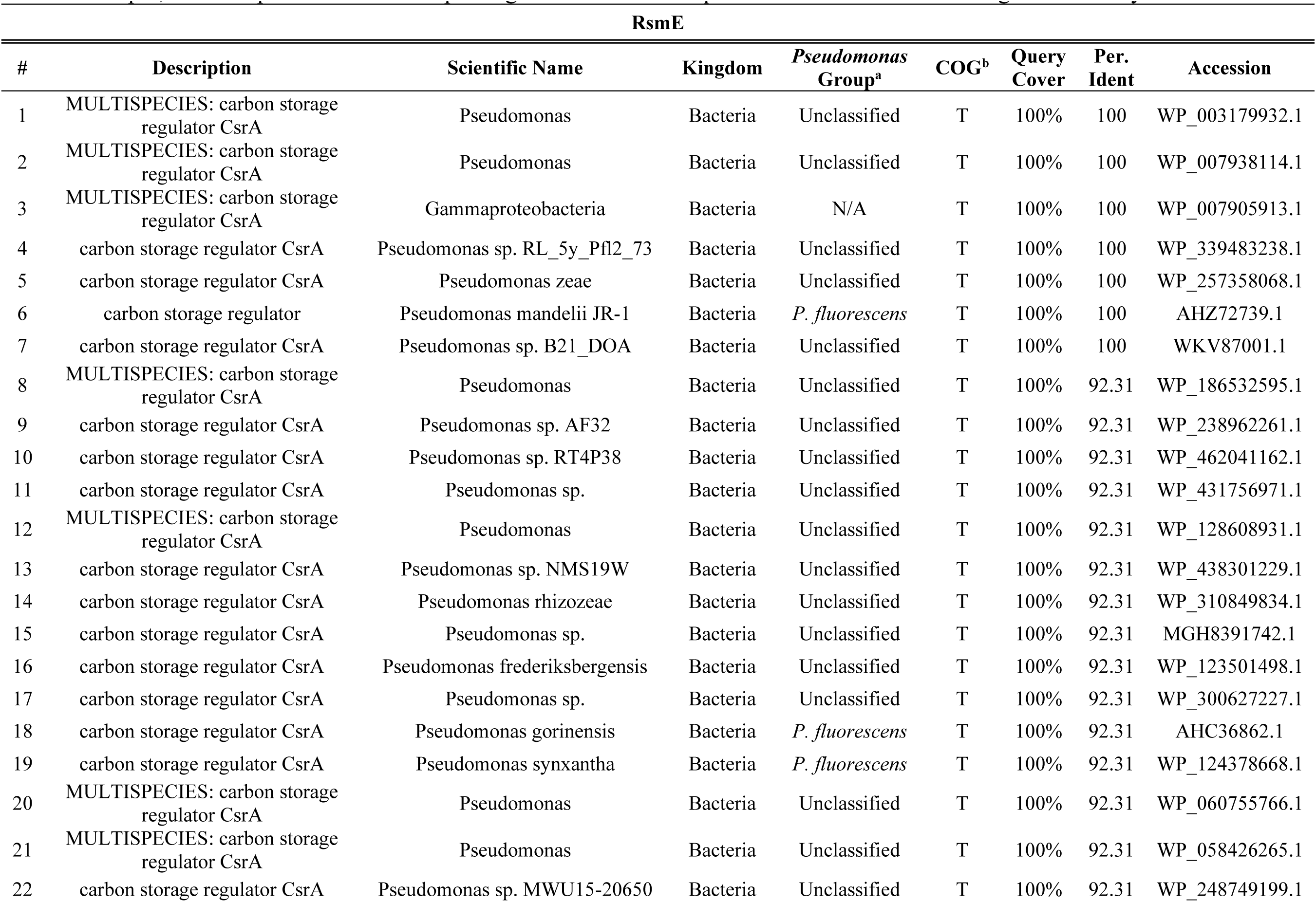
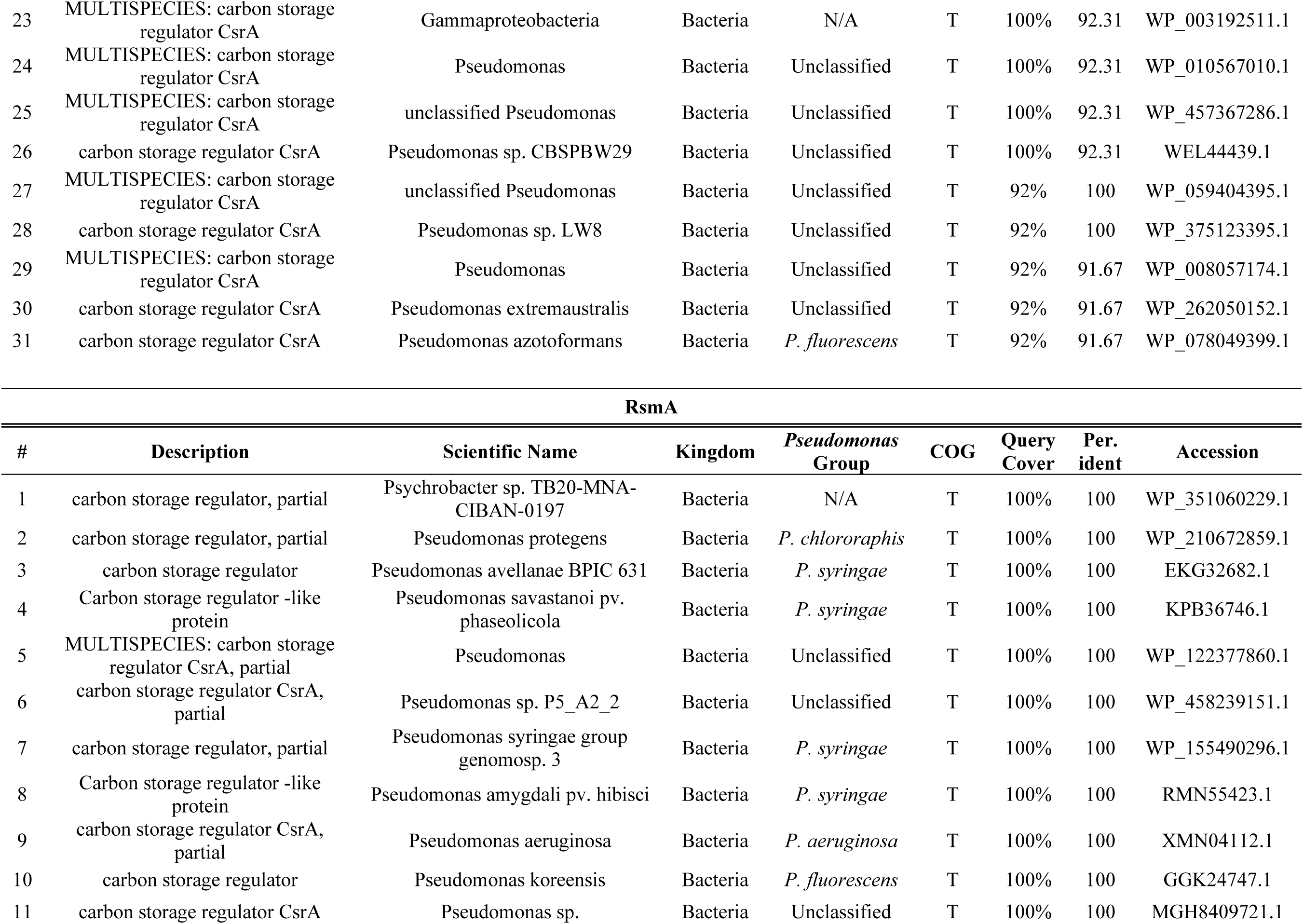
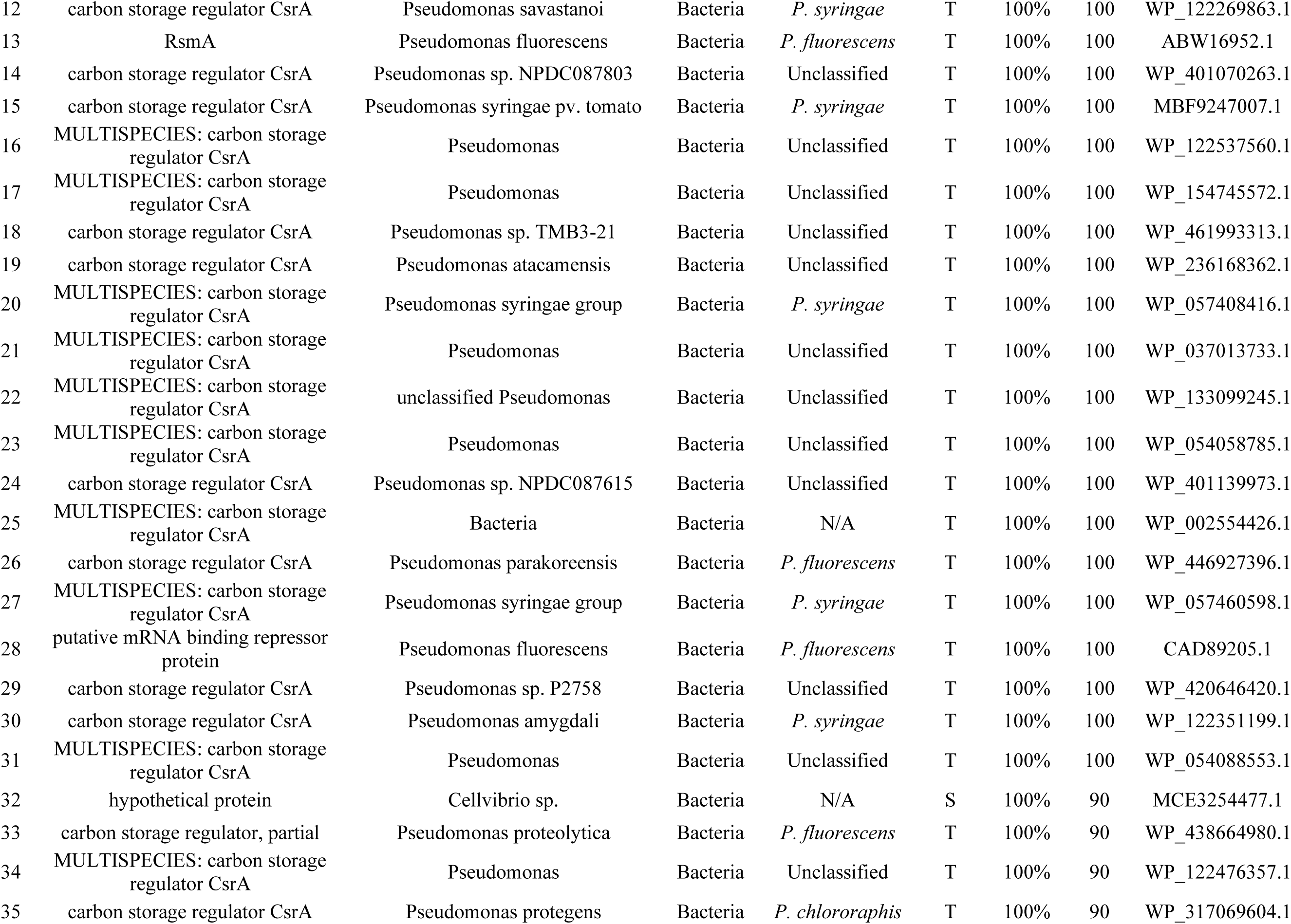
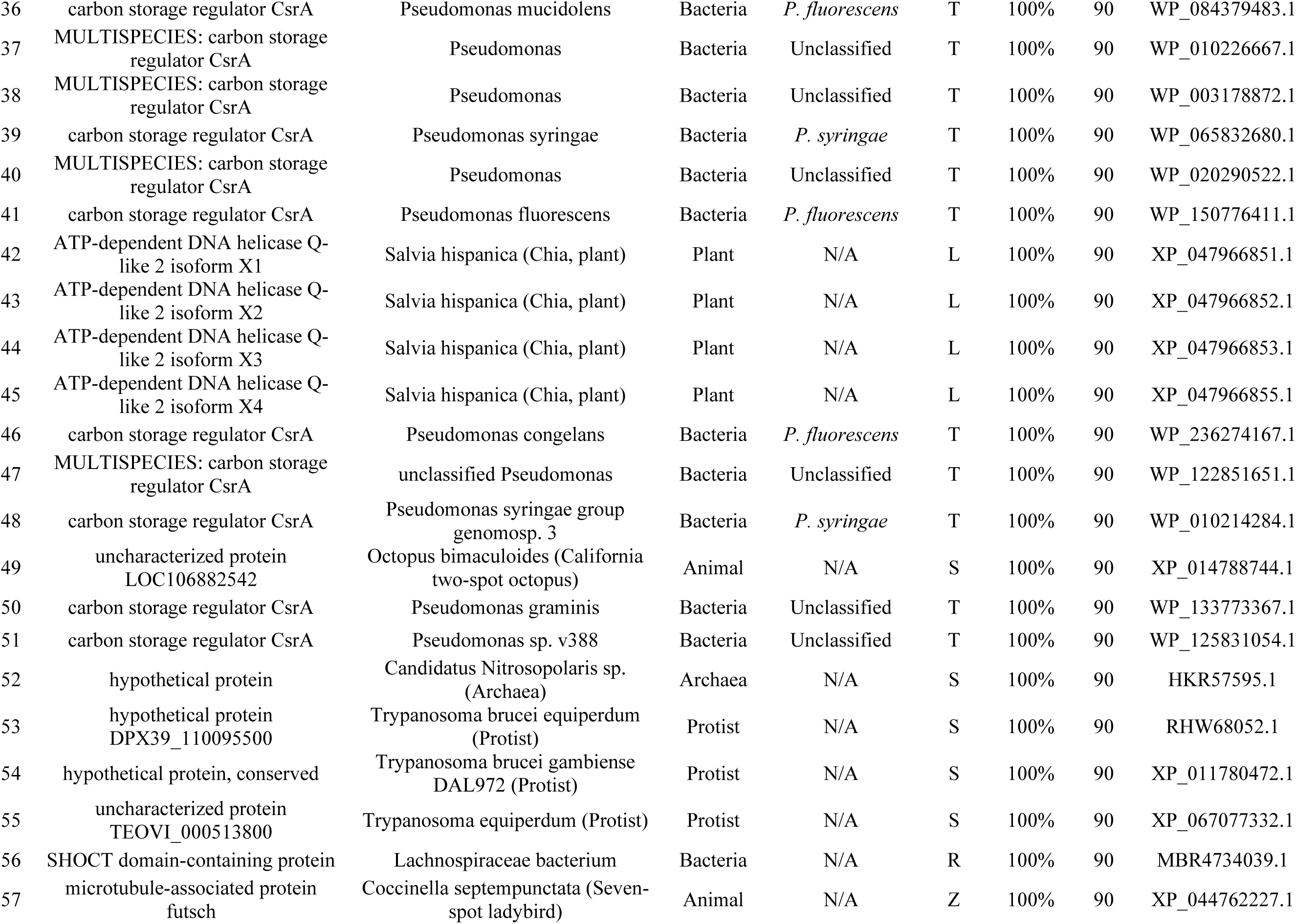
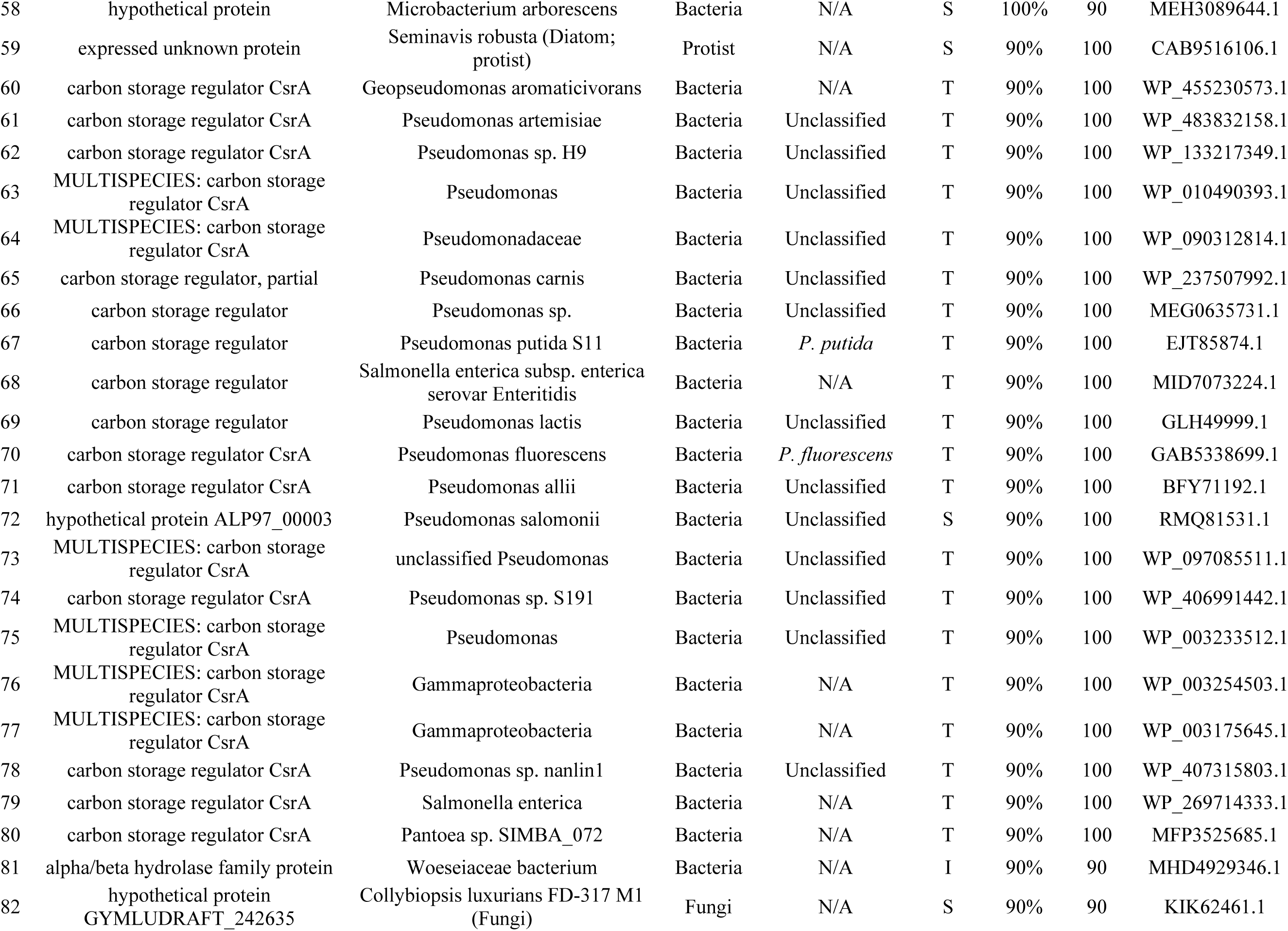
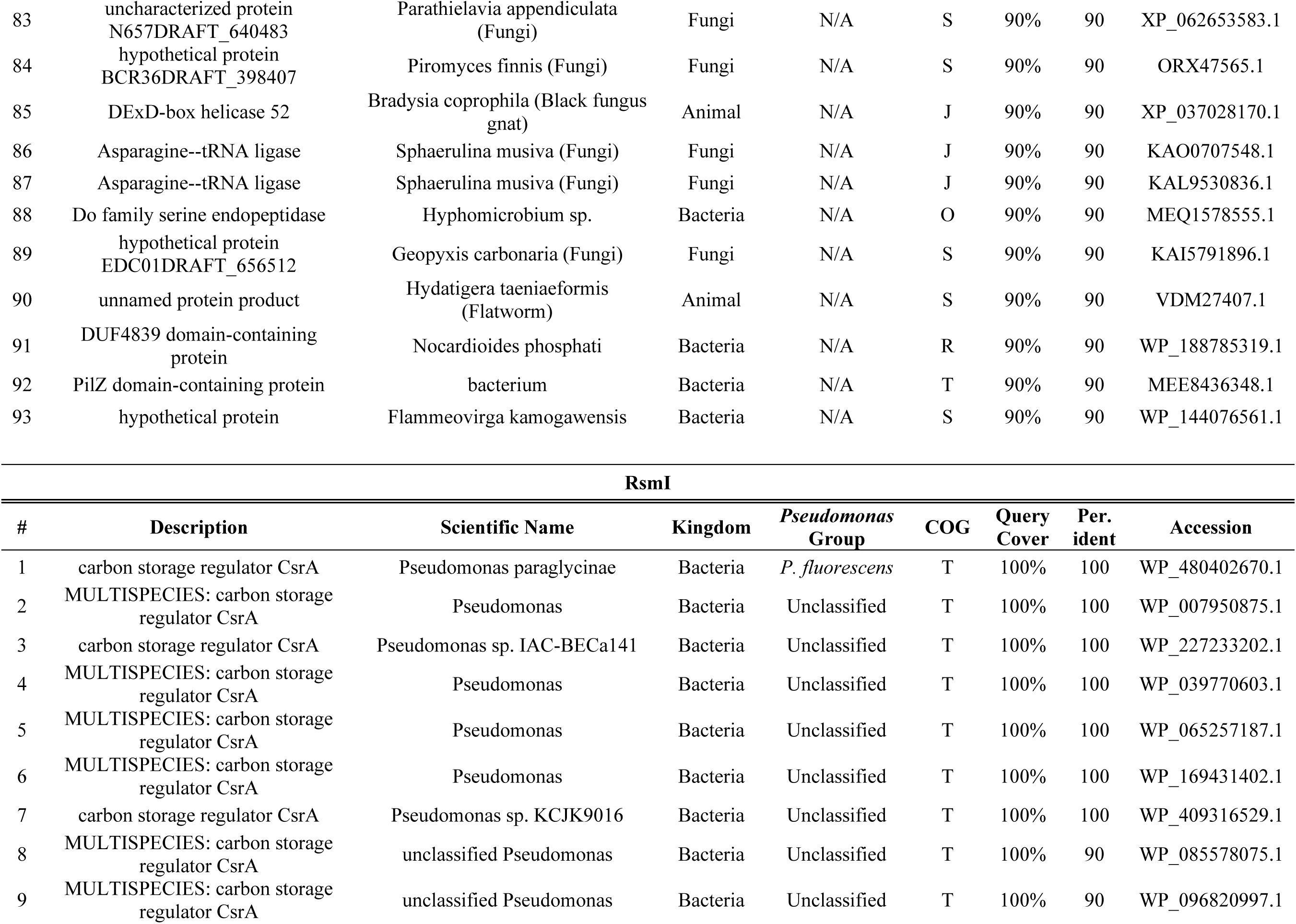
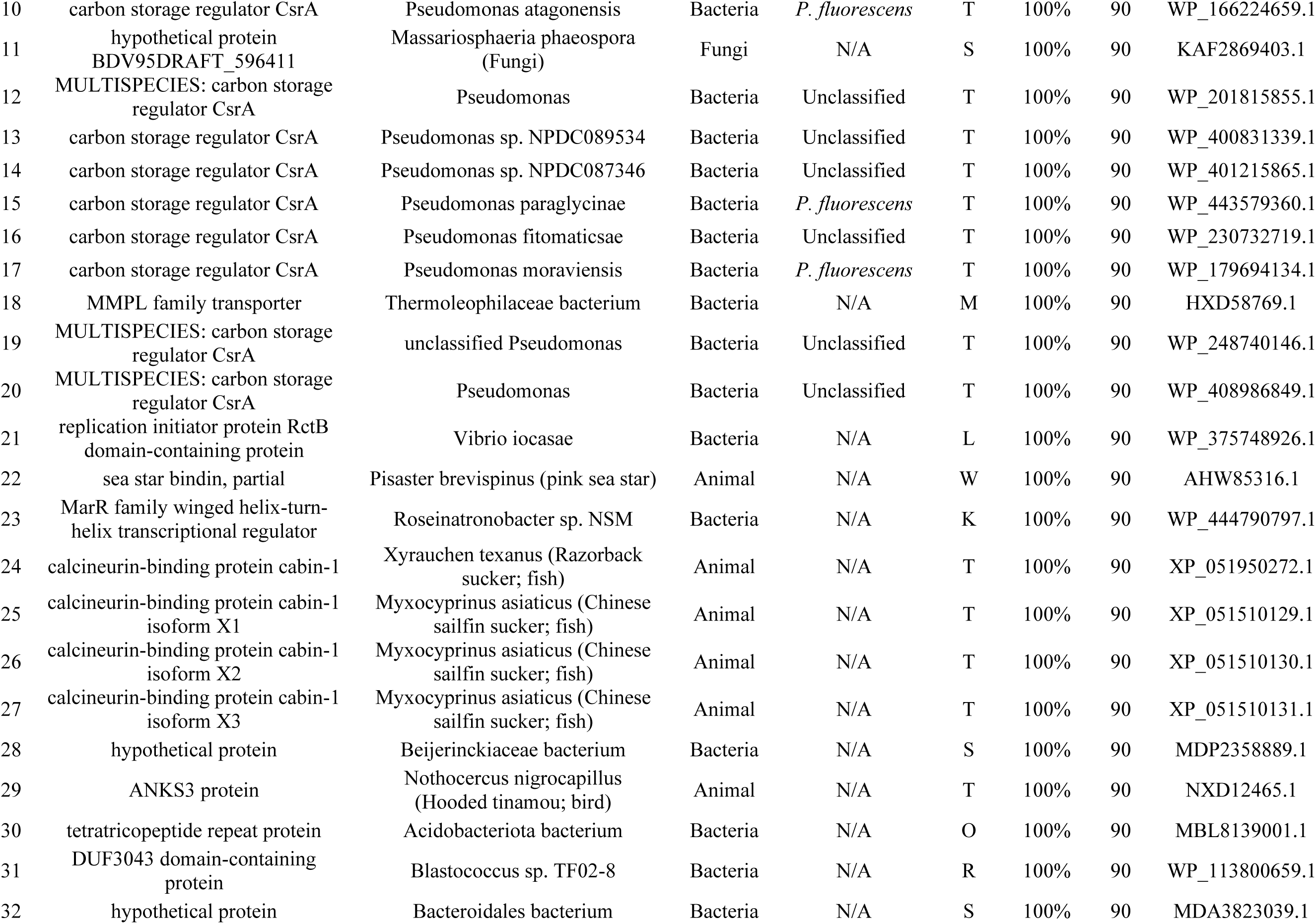
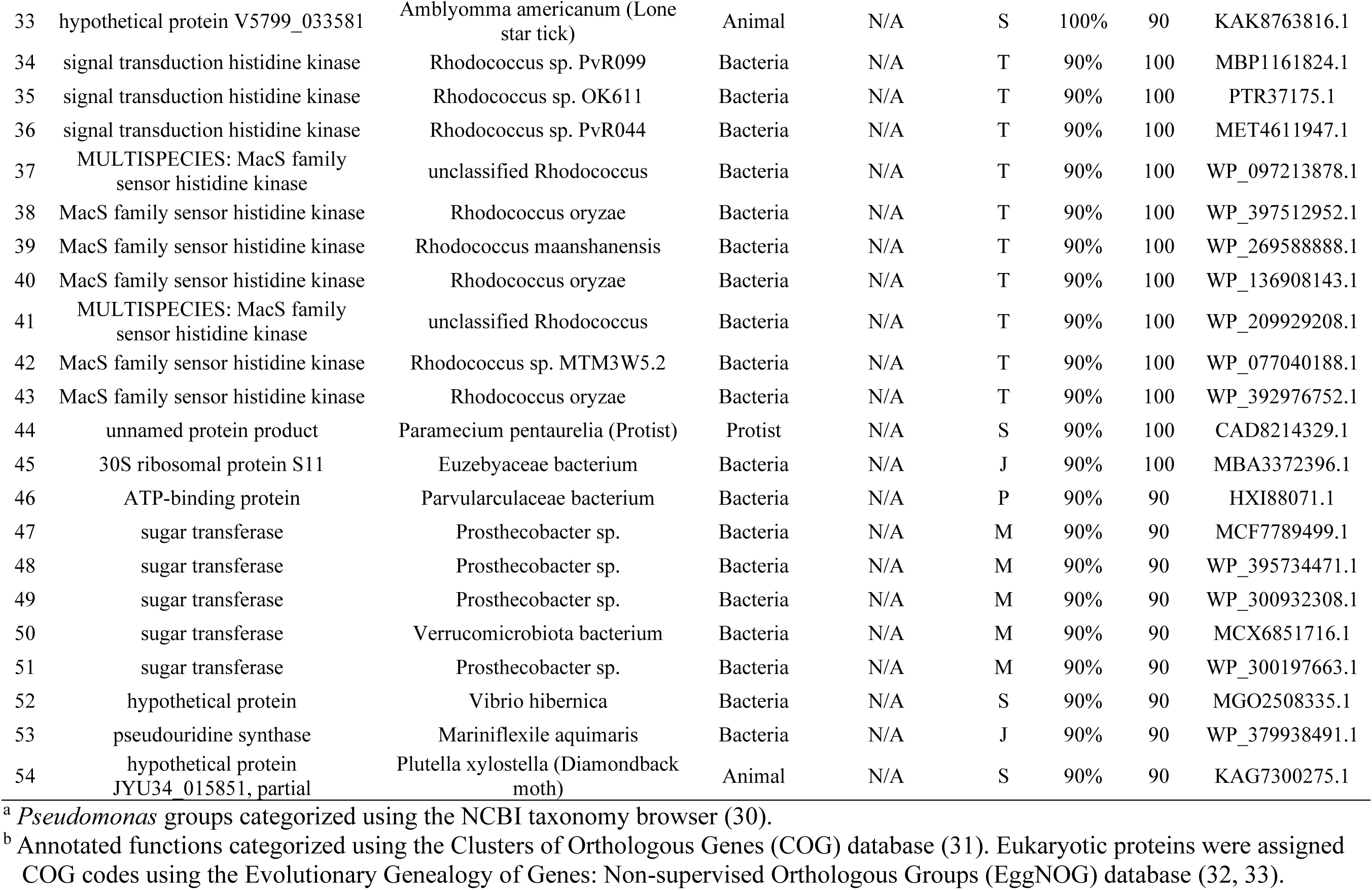
Top 1,000 Blastp hits of each Rsm paralog C-terminus tail sequence filtered for 90% coverage and identity.

| RsmE |  |  |  |  |  |  |  |  |
| --- | --- | --- | --- | --- | --- | --- | --- | --- |
| # | Description | Scientific Name | Kingdom | *Pseudomonas* Group<sup>a</sup> | COG<sup>b</sup> | Query Cover | Per. Ident | Accession |
| 1 | MULTISPECIES: carbon storage regulator CsrA | Pseudomonas | Bacteria | Unclassified | T | 100% | 100 | WP_003179932.1 |
| 2 | MULTISPECIES: carbon storage regulator CsrA | Pseudomonas | Bacteria | Unclassified | T | 100% | 100 | WP_007938114.1 |
| 3 | MULTISPECIES: carbon storage regulator CsrA | Gammaproteobacteria | Bacteria | N/A | T | 100% | 100 | WP_007905913.1 |
| 4 | carbon storage regulator CsrA | Pseudomonas sp. RL_5y_Pfl2_73 | Bacteria | Unclassified | T | 100% | 100 | WP_339483238.1 |
| 5 | carbon storage regulator CsrA | Pseudomonas zeae | Bacteria | Unclassified | T | 100% | 100 | WP_257358068.1 |
| 6 | carbon storage regulator | Pseudomonas mandelii JR-1 | Bacteria | *P. fluorescens* | T | 100% | 100 | AHZ72739.1 |
| 7 | carbon storage regulator CsrA | Pseudomonas sp. B21_DOA | Bacteria | Unclassified | T | 100% | 100 | WKV87001.1 |
| 8 | MULTISPECIES: carbon storage regulator CsrA | Pseudomonas | Bacteria | Unclassified | T | 100% | 92.31 | WP_186532595.1 |
| 9 | carbon storage regulator CsrA | Pseudomonas sp. AF32 | Bacteria | Unclassified | T | 100% | 92.31 | WP_238962261.1 |
| 10 | carbon storage regulator CsrA | Pseudomonas sp. RT4P38 | Bacteria | Unclassified | T | 100% | 92.31 | WP_462041162.1 |
| 11 | carbon storage regulator CsrA | Pseudomonas sp. | Bacteria | Unclassified | T | 100% | 92.31 | WP_431756971.1 |
| 12 | MULTISPECIES: carbon storage regulator CsrA | Pseudomonas | Bacteria | Unclassified | T | 100% | 92.31 | WP_128608931.1 |
| 13 | carbon storage regulator CsrA | Pseudomonas sp. NMS19W | Bacteria | Unclassified | T | 100% | 92.31 | WP_438301229.1 |
| 14 | carbon storage regulator CsrA | Pseudomonas rhizozeae | Bacteria | Unclassified | T | 100% | 92.31 | WP_310849834.1 |
| 15 | carbon storage regulator CsrA | Pseudomonas sp. | Bacteria | Unclassified | T | 100% | 92.31 | MGH8391742.1 |
| 16 | carbon storage regulator CsrA | Pseudomonas frederiksbergensis | Bacteria | Unclassified | T | 100% | 92.31 | WP_123501498.1 |
| 17 | carbon storage regulator CsrA | Pseudomonas sp. | Bacteria | Unclassified | T | 100% | 92.31 | WP_300627227.1 |
| 18 | carbon storage regulator CsrA | Pseudomonas gorinensis | Bacteria | *P. fluorescens* | T | 100% | 92.31 | AHC36862.1 |
| 19 | carbon storage regulator CsrA | Pseudomonas synxantha | Bacteria | *P. fluorescens* | T | 100% | 92.31 | WP_124378668.1 |
| 20 | MULTISPECIES: carbon storage regulator CsrA | Pseudomonas | Bacteria | Unclassified | T | 100% | 92.31 | WP_060755766.1 |
| 21 | MULTISPECIES: carbon storage regulator CsrA | Pseudomonas | Bacteria | Unclassified | T | 100% | 92.31 | WP_058426265.1 |
| 22 | carbon storage regulator CsrA | Pseudomonas sp. MWU15-20650 | Bacteria | Unclassified | T | 100% | 92.31 | WP_248749199.1 |
| 23 | MULTISPECIES: carbon storage regulator CsrA | Gammaproteobacteria | Bacteria | N/A | T | 100% | 92.31 | WP_003192511.1 |
| 24 | MULTISPECIES: carbon storage regulator CsrA | Pseudomonas | Bacteria | Unclassified | T | 100% | 92.31 | WP_010567010.1 |
| 25 | MULTISPECIES: carbon storage regulator CsrA | unclassified Pseudomonas | Bacteria | Unclassified | T | 100% | 92.31 | WP_457367286.1 |
| 26 | carbon storage regulator CsrA | Pseudomonas sp. CBSPBW29 | Bacteria | Unclassified | T | 100% | 92.31 | WEL44439.1 |
| 27 | MULTISPECIES: carbon storage regulator CsrA | unclassified Pseudomonas | Bacteria | Unclassified | T | 92% | 100 | WP_059404395.1 |
| 28 | carbon storage regulator CsrA | Pseudomonas sp. LW8 | Bacteria | Unclassified | T | 92% | 100 | WP_375123395.1 |
| 29 | MULTISPECIES: carbon storage regulator CsrA | Pseudomonas | Bacteria | Unclassified | T | 92% | 91.67 | WP_008057174.1 |
| 30 | carbon storage regulator CsrA | Pseudomonas extremaustralis | Bacteria | Unclassified | T | 92% | 91.67 | WP_262050152.1 |
| 31 | carbon storage regulator CsrA | Pseudomonas azotoformans | Bacteria | *P. fluorescens* | T | 92% | 91.67 | WP_078049399.1 |

RsmA
| # | Description | Scientific Name | Kingdom | *Pseudomonas* Group | COG | Query Cover | Per. ident | Accession |
| --- | --- | --- | --- | --- | --- | --- | --- | --- |
| 1 | carbon storage regulator, partial | Psychrobacter sp. TB20-MNA-CIBAN-0197 | Bacteria | N/A | T | 100% | 100 | WP_351060229.1 |
| 2 | carbon storage regulator, partial | Pseudomonas protegens | Bacteria | *P. chlororaphis* | T | 100% | 100 | WP_210672859.1 |
| 3 | carbon storage regulator | Pseudomonas avellanae BPIC 631 | Bacteria | *P. syringae* | T | 100% | 100 | EKG32682.1 |
| 4 | Carbon storage regulator -like protein | Pseudomonas savastanoi pv. phaseolicola | Bacteria | *P. syringae* | T | 100% | 100 | KPB36746.1 |
| 5 | MULTISPECIES: carbon storage regulator CsrA, partial | Pseudomonas | Bacteria | Unclassified | T | 100% | 100 | WP_122377860.1 |
| 6 | carbon storage regulator CsrA, partial | Pseudomonas sp. P5_A2_2 | Bacteria | Unclassified | T | 100% | 100 | WP_458239151.1 |
| 7 | carbon storage regulator, partial | Pseudomonas syringae group genomosp. 3 | Bacteria | *P. syringae* | T | 100% | 100 | WP_155490296.1 |
| 8 | Carbon storage regulator -like protein | Pseudomonas amygdali pv. hibisci | Bacteria | *P. syringae* | T | 100% | 100 | RMN55423.1 |
| 9 | carbon storage regulator CsrA, partial | Pseudomonas aeruginosa | Bacteria | *P. aeruginosa* | T | 100% | 100 | XMN04112.1 |
| 10 | carbon storage regulator | Pseudomonas koreensis | Bacteria | *P. fluorescens* | T | 100% | 100 | GGK24747.1 |
| 11 | carbon storage regulator CsrA | Pseudomonas sp. | Bacteria | Unclassified | T | 100% | 100 | MGH8409721.1 |
| 12 | carbon storage regulator CsrA | *Pseudomonas savastanoi* | Bacteria | *P. syringae* | T | 100% | 100 | WP_122269863.1 |
| 13 | RsmA | *Pseudomonas fluorescens* | Bacteria | *P. fluorescens* | T | 100% | 100 | ABW16952.1 |
| 14 | carbon storage regulator CsrA | *Pseudomonas* sp. NPDC087803 | Bacteria | Unclassified | T | 100% | 100 | WP_401070263.1 |
| 15 | carbon storage regulator CsrA | *Pseudomonas syringae* pv. tomato | Bacteria | *P. syringae* | T | 100% | 100 | MBF9247007.1 |
| 16 | MULTISPECIES: carbon storage regulator CsrA | *Pseudomonas* | Bacteria | Unclassified | T | 100% | 100 | WP_122537560.1 |
| 17 | MULTISPECIES: carbon storage regulator CsrA | *Pseudomonas* | Bacteria | Unclassified | T | 100% | 100 | WP_154745572.1 |
| 18 | carbon storage regulator CsrA | *Pseudomonas* sp. TMB3-21 | Bacteria | Unclassified | T | 100% | 100 | WP_461993313.1 |
| 19 | carbon storage regulator CsrA | *Pseudomonas atacamensis* | Bacteria | Unclassified | T | 100% | 100 | WP_236168362.1 |
| 20 | MULTISPECIES: carbon storage regulator CsrA | *Pseudomonas syringae* group | Bacteria | *P. syringae* | T | 100% | 100 | WP_057408416.1 |
| 21 | MULTISPECIES: carbon storage regulator CsrA | *Pseudomonas* | Bacteria | Unclassified | T | 100% | 100 | WP_037013733.1 |
| 22 | MULTISPECIES: carbon storage regulator CsrA | unclassified *Pseudomonas* | Bacteria | Unclassified | T | 100% | 100 | WP_133099245.1 |
| 23 | MULTISPECIES: carbon storage regulator CsrA | *Pseudomonas* | Bacteria | Unclassified | T | 100% | 100 | WP_054058785.1 |
| 24 | carbon storage regulator CsrA | *Pseudomonas* sp. NPDC087615 | Bacteria | Unclassified | T | 100% | 100 | WP_401139973.1 |
| 25 | MULTISPECIES: carbon storage regulator CsrA | Bacteria | Bacteria | N/A | T | 100% | 100 | WP_002554426.1 |
| 26 | carbon storage regulator CsrA | *Pseudomonas parakoreensis* | Bacteria | *P. fluorescens* | T | 100% | 100 | WP_446927396.1 |
| 27 | MULTISPECIES: carbon storage regulator CsrA | *Pseudomonas syringae* group | Bacteria | *P. syringae* | T | 100% | 100 | WP_057460598.1 |
| 28 | putative mRNA binding repressor protein | *Pseudomonas fluorescens* | Bacteria | *P. fluorescens* | T | 100% | 100 | CAD89205.1 |
| 29 | carbon storage regulator CsrA | *Pseudomonas* sp. P2758 | Bacteria | Unclassified | T | 100% | 100 | WP_420646420.1 |
| 30 | carbon storage regulator CsrA | *Pseudomonas amygdali* | Bacteria | *P. syringae* | T | 100% | 100 | WP_122351199.1 |
| 31 | MULTISPECIES: carbon storage regulator CsrA | *Pseudomonas* | Bacteria | Unclassified | T | 100% | 100 | WP_054088553.1 |
| 32 | hypothetical protein | *Cellvibrio* sp. | Bacteria | N/A | S | 100% | 90 | MCE3254477.1 |
| 33 | carbon storage regulator, partial | *Pseudomonas proteolytica* | Bacteria | *P. fluorescens* | T | 100% | 90 | WP_438664980.1 |
| 34 | MULTISPECIES: carbon storage regulator CsrA | *Pseudomonas* | Bacteria | Unclassified | T | 100% | 90 | WP_122476357.1 |
| 35 | carbon storage regulator CsrA | *Pseudomonas protegens* | Bacteria | *P. chlororaphis* | T | 100% | 90 | WP_317069604.1 |
| 36 | carbon storage regulator CsrA | *Pseudomonas mucidolens* | Bacteria | *P. fluorescens* | T | 100% | 90 | WP_084379483.1 |
| 37 | MULTISPECIES: carbon storage regulator CsrA | *Pseudomonas* | Bacteria | Unclassified | T | 100% | 90 | WP_010226667.1 |
| 38 | MULTISPECIES: carbon storage regulator CsrA | *Pseudomonas* | Bacteria | Unclassified | T | 100% | 90 | WP_003178872.1 |
| 39 | carbon storage regulator CsrA | *Pseudomonas syringae* | Bacteria | *P. syringae* | T | 100% | 90 | WP_065832680.1 |
| 40 | MULTISPECIES: carbon storage regulator CsrA | *Pseudomonas* | Bacteria | Unclassified | T | 100% | 90 | WP_020290522.1 |
| 41 | carbon storage regulator CsrA | *Pseudomonas fluorescens* | Bacteria | *P. fluorescens* | T | 100% | 90 | WP_150776411.1 |
| 42 | ATP-dependent DNA helicase Q-like 2 isoform X1 | *Salvia hispanica* (Chia, plant) | Plant | N/A | L | 100% | 90 | XP_047966851.1 |
| 43 | ATP-dependent DNA helicase Q-like 2 isoform X2 | *Salvia hispanica* (Chia, plant) | Plant | N/A | L | 100% | 90 | XP_047966852.1 |
| 44 | ATP-dependent DNA helicase Q-like 2 isoform X3 | *Salvia hispanica* (Chia, plant) | Plant | N/A | L | 100% | 90 | XP_047966853.1 |
| 45 | ATP-dependent DNA helicase Q-like 2 isoform X4 | *Salvia hispanica* (Chia, plant) | Plant | N/A | L | 100% | 90 | XP_047966855.1 |
| 46 | carbon storage regulator CsrA | *Pseudomonas congelans* | Bacteria | *P. fluorescens* | T | 100% | 90 | WP_236274167.1 |
| 47 | MULTISPECIES: carbon storage regulator CsrA | unclassified *Pseudomonas* | Bacteria | Unclassified | T | 100% | 90 | WP_122851651.1 |
| 48 | carbon storage regulator CsrA | *Pseudomonas syringae* group genomosp. 3 | Bacteria | *P. syringae* | T | 100% | 90 | WP_010214284.1 |
| 49 | uncharacterized protein LOC106882542 | *Octopus bimaculoides* (California two-spot octopus) | Animal | N/A | S | 100% | 90 | XP_014788744.1 |
| 50 | carbon storage regulator CsrA | *Pseudomonas graminis* | Bacteria | Unclassified | T | 100% | 90 | WP_133773367.1 |
| 51 | carbon storage regulator CsrA | *Pseudomonas* sp. v388 | Bacteria | Unclassified | T | 100% | 90 | WP_125831054.1 |
| 52 | hypothetical protein | *Candidatus Nitrosopolaris* sp. (Archaea) | Archaea | N/A | S | 100% | 90 | HKR57595.1 |
| 53 | hypothetical protein DPX39_110095500 | *Trypanosoma brucei equiperdum* (Protist) | Protist | N/A | S | 100% | 90 | RHW68052.1 |
| 54 | hypothetical protein, conserved | *Trypanosoma brucei gambiense* DAL972 (Protist) | Protist | N/A | S | 100% | 90 | XP_011780472.1 |
| 55 | uncharacterized protein TEOVI_000513800 | *Trypanosoma equiperdum* (Protist) | Protist | N/A | S | 100% | 90 | XP_067077332.1 |
| 56 | SHOCT domain-containing protein | Lachnospiraceae bacterium | Bacteria | N/A | R | 100% | 90 | MBR4734039.1 |
| 57 | microtubule-associated protein futsch | *Coccinella septempunctata* (Seven-spot ladybird) | Animal | N/A | Z | 100% | 90 | XP_044762227.1 |
| 58 | hypothetical protein | Microbacterium arborescens | Bacteria | N/A | S | 100% | 90 | MEH3089644.1 |
| 59 | expressed unknown protein | Seminavis robusta (Diatom; protist) | Protist | N/A | S | 90% | 100 | CAB9516106.1 |
| 60 | carbon storage regulator CsrA | Geopseudomonas aromaticivorans | Bacteria | N/A | T | 90% | 100 | WP_455230573.1 |
| 61 | carbon storage regulator CsrA | Pseudomonas artemisiae | Bacteria | Unclassified | T | 90% | 100 | WP_483832158.1 |
| 62 | carbon storage regulator CsrA | Pseudomonas sp. H9 | Bacteria | Unclassified | T | 90% | 100 | WP_133217349.1 |
| 63 | MULTISPECIES: carbon storage regulator CsrA | Pseudomonas | Bacteria | Unclassified | T | 90% | 100 | WP_010490393.1 |
| 64 | MULTISPECIES: carbon storage regulator CsrA | Pseudomonadaceae | Bacteria | Unclassified | T | 90% | 100 | WP_090312814.1 |
| 65 | carbon storage regulator, partial | Pseudomonas carnis | Bacteria | Unclassified | T | 90% | 100 | WP_237507992.1 |
| 66 | carbon storage regulator | Pseudomonas sp. | Bacteria | Unclassified | T | 90% | 100 | MEG0635731.1 |
| 67 | carbon storage regulator | Pseudomonas putida S11 | Bacteria | *P. putida* | T | 90% | 100 | EJT85874.1 |
| 68 | carbon storage regulator | Salmonella enterica subsp. enterica serovar Enteritidis | Bacteria | N/A | T | 90% | 100 | MID7073224.1 |
| 69 | carbon storage regulator | Pseudomonas lactis | Bacteria | Unclassified | T | 90% | 100 | GLH49999.1 |
| 70 | carbon storage regulator CsrA | Pseudomonas fluorescens | Bacteria | *P. fluorescens* | T | 90% | 100 | GAB5338699.1 |
| 71 | carbon storage regulator CsrA | Pseudomonas allii | Bacteria | Unclassified | T | 90% | 100 | BFY71192.1 |
| 72 | hypothetical protein ALP97_00003 | Pseudomonas salomonii | Bacteria | Unclassified | S | 90% | 100 | RMQ81531.1 |
| 73 | MULTISPECIES: carbon storage regulator CsrA | unclassified Pseudomonas | Bacteria | Unclassified | T | 90% | 100 | WP_097085511.1 |
| 74 | carbon storage regulator CsrA | Pseudomonas sp. S191 | Bacteria | Unclassified | T | 90% | 100 | WP_406991442.1 |
| 75 | MULTISPECIES: carbon storage regulator CsrA | Pseudomonas | Bacteria | Unclassified | T | 90% | 100 | WP_003233512.1 |
| 76 | MULTISPECIES: carbon storage regulator CsrA | Gammaproteobacteria | Bacteria | N/A | T | 90% | 100 | WP_003254503.1 |
| 77 | MULTISPECIES: carbon storage regulator CsrA | Gammaproteobacteria | Bacteria | N/A | T | 90% | 100 | WP_003175645.1 |
| 78 | carbon storage regulator CsrA | Pseudomonas sp. nanlin1 | Bacteria | Unclassified | T | 90% | 100 | WP_407315803.1 |
| 79 | carbon storage regulator CsrA | Salmonella enterica | Bacteria | N/A | T | 90% | 100 | WP_269714333.1 |
| 80 | carbon storage regulator CsrA | Pantoea sp. SIMBA_072 | Bacteria | N/A | T | 90% | 100 | MFP3525685.1 |
| 81 | alpha/beta hydrolase family protein | Woeseiaceae bacterium | Bacteria | N/A | I | 90% | 90 | MHD4929346.1 |
| 82 | hypothetical protein GYMLUDRAFT_242635 | Collybiopsis luxurians FD-317 M1 (Fungi) | Fungi | N/A | S | 90% | 90 | KIK62461.1 |
| 83 | uncharacterized protein<br>N657DRAFT_640483 | Parathielavia appendiculata<br>(Fungi) | Fungi | N/A | S | 90% | 90 | XP_062653583.1 |
| 84 | hypothetical protein<br>BCR36DRAFT_398407 | Piromyces finnis (Fungi) | Fungi | N/A | S | 90% | 90 | ORX47565.1 |
| 85 | DExD-box helicase 52 | Bradysia coprophila (Black fungus<br>gnat) | Animal | N/A | J | 90% | 90 | XP_037028170.1 |
| 86 | Asparagine--tRNA ligase | Sphaerulina musiva (Fungi) | Fungi | N/A | J | 90% | 90 | KAO0707548.1 |
| 87 | Asparagine--tRNA ligase | Sphaerulina musiva (Fungi) | Fungi | N/A | J | 90% | 90 | KAL9530836.1 |
| 88 | Do family serine endopeptidase | Hyphomicrobium sp. | Bacteria | N/A | O | 90% | 90 | MEQ1578555.1 |
| 89 | hypothetical protein<br>EDC01DRAFT_656512 | Geopyxis carbonaria (Fungi) | Fungi | N/A | S | 90% | 90 | KAI5791896.1 |
| 90 | unnamed protein product | Hydatigera taeniaeformis<br>(Flatworm) | Animal | N/A | S | 90% | 90 | VDM27407.1 |
| 91 | DUF4839 domain-containing<br>protein | Nocardioides phosphati | Bacteria | N/A | R | 90% | 90 | WP_188785319.1 |
| 92 | PilZ domain-containing protein | bacterium | Bacteria | N/A | T | 90% | 90 | MEE8436348.1 |
| 93 | hypothetical protein | Flammeovirga kamogawensis | Bacteria | N/A | S | 90% | 90 | WP_144076561.1 |

RsmI
| # | Description | Scientific Name | Kingdom | *Pseudomonas*<br>Group | COG | Query<br>Cover | Per.<br>ident | Accession |
| --- | --- | --- | --- | --- | --- | --- | --- | --- |
| 1 | carbon storage regulator CsrA | Pseudomonas paraglycinae | Bacteria | *P. fluorescens* | T | 100% | 100 | WP_480402670.1 |
| 2 | MULTISPECIES: carbon storage<br>regulator CsrA | Pseudomonas | Bacteria | Unclassified | T | 100% | 100 | WP_007950875.1 |
| 3 | carbon storage regulator CsrA | Pseudomonas sp. IAC-BECa141 | Bacteria | Unclassified | T | 100% | 100 | WP_227233202.1 |
| 4 | MULTISPECIES: carbon storage<br>regulator CsrA | Pseudomonas | Bacteria | Unclassified | T | 100% | 100 | WP_039770603.1 |
| 5 | MULTISPECIES: carbon storage<br>regulator CsrA | Pseudomonas | Bacteria | Unclassified | T | 100% | 100 | WP_065257187.1 |
| 6 | MULTISPECIES: carbon storage<br>regulator CsrA | Pseudomonas | Bacteria | Unclassified | T | 100% | 100 | WP_169431402.1 |
| 7 | carbon storage regulator CsrA | Pseudomonas sp. KCJK9016 | Bacteria | Unclassified | T | 100% | 100 | WP_409316529.1 |
| 8 | MULTISPECIES: carbon storage<br>regulator CsrA | unclassified Pseudomonas | Bacteria | Unclassified | T | 100% | 90 | WP_085578075.1 |
| 9 | MULTISPECIES: carbon storage<br>regulator CsrA | unclassified Pseudomonas | Bacteria | Unclassified | T | 100% | 90 | WP_096820997.1 |
| 10 | carbon storage regulator CsrA | *Pseudomonas atagonensis* | Bacteria | *P. fluorescens* | T | 100% | 90 | WP_166224659.1 |
| 11 | hypothetical protein<br>BDV95DRAFT_596411 | *Massariosphaeria phaeospora*<br>(Fungi) | Fungi | N/A | S | 100% | 90 | KAF2869403.1 |
| 12 | MULTISPECIES: carbon storage<br>regulator CsrA | *Pseudomonas* | Bacteria | Unclassified | T | 100% | 90 | WP_201815855.1 |
| 13 | carbon storage regulator CsrA | *Pseudomonas* sp. NPDC089534 | Bacteria | Unclassified | T | 100% | 90 | WP_400831339.1 |
| 14 | carbon storage regulator CsrA | *Pseudomonas* sp. NPDC087346 | Bacteria | Unclassified | T | 100% | 90 | WP_401215865.1 |
| 15 | carbon storage regulator CsrA | *Pseudomonas paraglycinae* | Bacteria | *P. fluorescens* | T | 100% | 90 | WP_443579360.1 |
| 16 | carbon storage regulator CsrA | *Pseudomonas fitomaticsae* | Bacteria | Unclassified | T | 100% | 90 | WP_230732719.1 |
| 17 | carbon storage regulator CsrA | *Pseudomonas moraviensis* | Bacteria | *P. fluorescens* | T | 100% | 90 | WP_179694134.1 |
| 18 | MMPL family transporter | Thermoleophilaceae bacterium | Bacteria | N/A | M | 100% | 90 | HXD58769.1 |
| 19 | MULTISPECIES: carbon storage<br>regulator CsrA | unclassified *Pseudomonas* | Bacteria | Unclassified | T | 100% | 90 | WP_248740146.1 |
| 20 | MULTISPECIES: carbon storage<br>regulator CsrA | *Pseudomonas* | Bacteria | Unclassified | T | 100% | 90 | WP_408986849.1 |
| 21 | replication initiator protein RctB<br>domain-containing protein | *Vibrio iocasae* | Bacteria | N/A | L | 100% | 90 | WP_375748926.1 |
| 22 | sea star bindin, partial | *Pisaster brevispinus* (pink sea star) | Animal | N/A | W | 100% | 90 | AHW85316.1 |
| 23 | MarR family winged helix-turn-<br>helix transcriptional regulator | *Roseinatronobacter* sp. NSM | Bacteria | N/A | K | 100% | 90 | WP_444790797.1 |
| 24 | calcineurin-binding protein cabin-1 | *Xyrauchen texanus* (Razorback<br>sucker; fish) | Animal | N/A | T | 100% | 90 | XP_051950272.1 |
| 25 | calcineurin-binding protein cabin-1<br>isoform X1 | *Myxocyprinus asiaticus* (Chinese<br>sailfin sucker; fish) | Animal | N/A | T | 100% | 90 | XP_051510129.1 |
| 26 | calcineurin-binding protein cabin-1<br>isoform X2 | *Myxocyprinus asiaticus* (Chinese<br>sailfin sucker; fish) | Animal | N/A | T | 100% | 90 | XP_051510130.1 |
| 27 | calcineurin-binding protein cabin-1<br>isoform X3 | *Myxocyprinus asiaticus* (Chinese<br>sailfin sucker; fish) | Animal | N/A | T | 100% | 90 | XP_051510131.1 |
| 28 | hypothetical protein | Beijerinckiaceae bacterium | Bacteria | N/A | S | 100% | 90 | MDP2358889.1 |
| 29 | ANKS3 protein | *Nothocercus nigrocapillus*<br>(Hooded tinamou; bird) | Animal | N/A | T | 100% | 90 | NXD12465.1 |
| 30 | tetratricopeptide repeat protein | *Acidobacteriota* bacterium | Bacteria | N/A | O | 100% | 90 | MBL8139001.1 |
| 31 | DUF3043 domain-containing<br>protein | *Blastococcus* sp. TF02-8 | Bacteria | N/A | R | 100% | 90 | WP_113800659.1 |
| 32 | hypothetical protein | Bacteroidales bacterium | Bacteria | N/A | S | 100% | 90 | MDA3823039.1 |
| 33 | hypothetical protein V5799_033581 | Amblyomma americanum (Lone star tick) | Animal | N/A | S | 100% | 90 | KAK8763816.1 |
| 34 | signal transduction histidine kinase | Rhodococcus sp. PvR099 | Bacteria | N/A | T | 90% | 100 | MBP1161824.1 |
| 35 | signal transduction histidine kinase | Rhodococcus sp. OK611 | Bacteria | N/A | T | 90% | 100 | PTR37175.1 |
| 36 | signal transduction histidine kinase | Rhodococcus sp. PvR044 | Bacteria | N/A | T | 90% | 100 | MET4611947.1 |
| 37 | MULTISPECIES: MacS family sensor histidine kinase | unclassified Rhodococcus | Bacteria | N/A | T | 90% | 100 | WP_097213878.1 |
| 38 | MacS family sensor histidine kinase | Rhodococcus oryzae | Bacteria | N/A | T | 90% | 100 | WP_397512952.1 |
| 39 | MacS family sensor histidine kinase | Rhodococcus maanshanensis | Bacteria | N/A | T | 90% | 100 | WP_269588888.1 |
| 40 | MacS family sensor histidine kinase | Rhodococcus oryzae | Bacteria | N/A | T | 90% | 100 | WP_136908143.1 |
| 41 | MULTISPECIES: MacS family sensor histidine kinase | unclassified Rhodococcus | Bacteria | N/A | T | 90% | 100 | WP_209929208.1 |
| 42 | MacS family sensor histidine kinase | Rhodococcus sp. MTM3W5.2 | Bacteria | N/A | T | 90% | 100 | WP_077040188.1 |
| 43 | MacS family sensor histidine kinase | Rhodococcus oryzae | Bacteria | N/A | T | 90% | 100 | WP_392976752.1 |
| 44 | unnamed protein product | Paramecium pentaurelia (Protist) | Protist | N/A | S | 90% | 100 | CAD8214329.1 |
| 45 | 30S ribosomal protein S11 | Euzebyaceae bacterium | Bacteria | N/A | J | 90% | 100 | MBA3372396.1 |
| 46 | ATP-binding protein | Parvularculaceae bacterium | Bacteria | N/A | P | 90% | 90 | HXI88071.1 |
| 47 | sugar transferase | Prostheco bacter sp. | Bacteria | N/A | M | 90% | 90 | MCF7789499.1 |
| 48 | sugar transferase | Prostheco bacter sp. | Bacteria | N/A | M | 90% | 90 | WP_395734471.1 |
| 49 | sugar transferase | Prostheco bacter sp. | Bacteria | N/A | M | 90% | 90 | WP_300932308.1 |
| 50 | sugar transferase | Verrucomicrobiota bacterium | Bacteria | N/A | M | 90% | 90 | MCX6851716.1 |
| 51 | sugar transferase | Prostheco bacter sp. | Bacteria | N/A | M | 90% | 90 | WP_300197663.1 |
| 52 | hypothetical protein | Vibrio hibernica | Bacteria | N/A | S | 90% | 90 | MGO2508335.1 |
| 53 | pseudouridine synthase | Mariniflexile aquimaris | Bacteria | N/A | J | 90% | 90 | WP_379938491.1 |
| 54 | hypothetical protein JYU34_015851, partial | Plutella xylostella (Diamondback moth) | Animal | N/A | S | 90% | 90 | KAG7300275.1 |

<sup>a</sup> *Pseudomonas* groups categorized using the NCBI taxonomy browser (30).
<sup>b</sup> Annotated functions categorized using the Clusters of Orthologous Genes (COG) database (31). Eukaryotic proteins were assigned COG codes using the Evolutionary Genealogy of Genes: Non-supervised Orthologous Groups (EggNOG) database (32, 33).

